# Evolutionary insights into PtdIns3*P* signaling through FYVE and PHOX effector proteins from the moss *Physcomitrella patens*

**DOI:** 10.1101/765594

**Authors:** Patricia Agudelo-Romero, Ana Margarida Fortes, Trinidad Suárez, Hernán Ramiro Lascano, Laura Saavedra

## Abstract

Phosphatidylinositol 3-phosphate (PtdIns3*P*) is one of the five different phosphoinositides (PPIs) species in plant cells, which regulate several aspects of plant growth and development, as well as responses to biotic and abiotic stresses. The mechanistic insights underlying PtdIns3*P* mode of action, specifically through PtdIns3*P*-binding effectors such as FYVE and PHOX proteins have been partially explored in plants with main focus on *Arabidopsis thaliana.* Additionally, they have been underexplored in other plant organisms such as bryophytes, the earliest diverging group of terrestrial flora.

In this study, we searched for genes coding for FYVE and PHOX domains containing sequences from different photosynthetic organisms in order to gather evolutionary insights on these PPI binding domains, followed by an *in silico* characterization of the *FYVE* and *PHOX* gene family in the moss *Physcomitrella patens*. Phylogenetic analysis showed that PpFYVE proteins can be grouped in 7 subclasses, with an additional subclass whose FYVE domain was lost during evolution to higher plants. On the other hand, PpPHOX proteins are classified into 5 subclasses. Expression analyses based on RNAseq data together with the analysis of *cis*-acting regulatory elements and transcription factor binding sites in promoter regions suggest the importance of these proteins in regulating stress responses but mainly developmental processes in *P. patens*. The results provide valuable information and robust candidate genes for future functional analysis aiming to further explore the role of this signaling pathway mainly during growth and development of tip growing cells and during the transition from 2D to 3D growth, which could provide ancestral regulatory players undertaken during plant evolution.

## INTRODUCTION

Phosphoinositides (PPIs) are eukaryotic signaling phospholipids derivatives of phosphatidylinositol (PI), which control a large number of cellular functions through their role as regulators of membrane dynamics (Schink, Tan, and Stenmark 2016). In plants, PPIs control central aspects of plant growth and development, as well as responses to biotic and abiotic stresses (Gerth et al. 2017). Severe phenotypes have been described using an overexpression or a down-regulation approach for several members of the plant PPI-metabolism enzyme network. However, this approach can lead to misleading interpretations of resulting phenotypes and associated PPI-specific biological functions because affecting one member of this network frequently leads to the adjustment of other members of PPI-metabolism, in addition to the fact that, some PPI species exist in multiple locations performing different functions.

One of the mechanisms by which PPIs control cellular processes is by their ability to recruit specific phosphoinositide binding protein effectors to membranes in a reversible manner achieved by both, a low to moderate affinity of protein effectors for PPIs, and by PPIs turnover by phosphorylation or dephosphorylation (Moravcevic, Oxley, and Lemmon 2012). Therefore, addressing the role of effectors and downstream targets of PPI regulation will improve our knowledge on PPI-specific functions in plants.

Phosphatidylinositol 3-phosphate (PtdIns3*P*) is one of the five different PPI species identified in plant cells and focus of this study. Protein effectors containing a PtdIns3*P*-specific binding module are denoted as FYVE and PHOX proteins. FYVE domains, named for the four proteins in which were first identified (<u>F</u>ab1, <u>Y</u>OTB, <u>V</u>ac1, <u>E</u>EA) are zinc binding fingers containing ∼70 amino acids, comprising two double-stranded antiparallel β-sheets and a small C-terminal α-helix that are held together by two tetrahedrally-coordinated Zn^2+^ ions (Kutateladze 2006). This domain is defined by three conserved sequences that form a highly positively charged binding site for PtdIns3*P*: the WxxD, (R/K)(R/K)HHCR and RVC motifs (Kutateladze 2006). Upon binding of the FYVE domain to PtdIns3*P* containing membranes, FYVE domains are able to penetrate into the bilayers through an exposed hydrophobic protrusion known as the membrane insertion loop (MIL). The PHOX domain (also known as PX) was originally named for its presence in the p40^phox^ and p47^phox^ subunits of the phagocyte NADPH oxidase complex. Although most PHOX domains bind to PtdIns3*P*, affinities for other PPIs and anionic lipids have also been reported. A recently thorough description of the membrane-binding specificities of the human PX domain family resolve four distinct categories of PX domains; including binding specifically to PtdIns3*P*, binding other PPIs as well as PtdIns3*P*, binding other lipids but not PtdIns3*P*, and those that do not appear to bind membrane lipids significantly at all (Chandra et al. 2019). PHOX domains comprise a region of 130 amino acids with a relative weak primary sequence identity but substantial similarity at the level of the three dimensional structure. The overall structure consists of a PI-binding pocket including basic residues, three β-strands followed by three α-helices, and a proline-rich loop with the conserved sequence YPxxPxK (Y denotes large aliphatic amino acids V,I,L,M) between the first and second helix (Seet and Hong 2006). In most cases, the FYVE and PHOX domains bind weakly to PtdIns3*P* and membrane assembly appear to rely additionally on effector oligomerization to increase avidity of membrane binding, or coincident detection of another membrane molecule, membrane charge or membrane curvature (Schink, Raiborg, and Stenmark 2013).

*In silico* genome wide analyses have predicted the presence of 15 and 19 FYVE proteins in *A. thaliana* and *Oryza sativa*, respectively, and 11 PHOX members in both species (Xiao and Shaw 2016, Wywial and Singh 2010, van Leeuwen et al. 2004, Jensen et al. 2001). Up to date, a few FYVE and PHOX proteins have been functionally characterized, mainly in *A. thaliana*, revealing that PtdIns3*P* is a crucial player in endocytic trafficking, vesicular transport and regulation of autophagy. An example of FYVE-domain proteins are FAB enzymes, which synthesize PtdIns(3,5)*P*_2_ from PtdIns3*P*, and are essential for endomembrane homeostasis including endocytosis, vacuole formation and vacuolar acidification in *A. thaliana* (Hirano et al. 2011, Hirano, Munnik, and Sato 2015, 2017, Serrazina, Dias, and Malhó 2014, Bak et al. 2013, Whitley, Hinz, and Doughty 2009). FYVE1/FREE1, another FYVE-domain protein, was shown to participate in several processes such as the regulation of the formation of intraluminal vesicles (ILVs) in prevacuolar compartments/multivesicular bodies (PVCs/MVBs), MVB-mediated sorting and degradation of ubiquitylated membrane proteins, vacuolar protein transport, autophagic degradation, and vacuole biogenesis (Gao et al. 2014, Gao et al. 2015, Barberon et al. 2014, Kolb et al. 2015, Belda-Palazon et al. 2016).

In the case of PHOX domain proteins, the syntaxin family has been the more extensively studied. AtSNX1 function as a sorting endosome from which endocytosed plasma membrane (PM) proteins, such as the Iron-Regulated Transporter1 (IRT1) or the auxin efflux carrier PIN2 and secretory proteins are sorted to diverse destinations (Ivanov et al. 2014, Jaillais et al. 2008, Jaillais et al. 2006, Pourcher et al. 2010). In addition, AtSNX2a and AtSNX2b, have distinct functions in trafficking of storage proteins and plant development (Pourcher et al. 2010), whereas AtEREX, another PHOX-domain protein, was recently suggested as a genuine effector of canonical RAB5s in *A. thaliana* (Sakurai et al. 2016).

As more species have their complete reference genome sequenced, additional *FYVE* and *PHOX* genes can be identified and reveal the biological roles of these gene families. Due to the fact that most available genomes are from flowering plants, covering other rich-species lineages, in particular bryophytes-comprising mosses, liverworts and hornworts-may reveal novel evolutionary paths/functions that have been undertaken during evolution (Rensing et al. 2008, Rensing 2017). The moss *Physcomitrella patens* is a powerful model system for evolutionary and functional genomics in plants (Rensing et al. 2008, Collonnier et al. 2016, Schaefer and Zrÿd 1997, Bezanilla, Pan, and Quatrano 2003). It belongs to the bryophytes, the earliest diverging group of terrestrial flora. Bryophytes constitute a key group to understand the genetic basis of the critical innovations that allowed green plants to evolve from an aquatic ancestor and to adapt to the terrestrial environment (Rensing et al. 2008). In addition, its relatively simple body architecture, an haploid-dominant life cycle, excellent cytology combined with the fact that protonemata and rhizoid cells exhibit polar growth, provide the opportunity to study signal transduction mechanisms at a molecular level, particularly appropriated for processes where vesicle trafficking and secretion are involved (Vidali and Bezanilla 2012, Cove et al. 2006).

We have previously analyzed ^32^P-radiolabeled phospholipids extracted from a suspension culture of *P. patens* protonemal tissue under normal growth conditions and identified a similar PPI composition to that of flowering plants (Saavedra et al. 2009). PtdIns3*P* and PtdIns(3,5)*P*_2_ are minor PPIs in *P. patens*, which closely agrees with what is observed in other organisms (Saavedra et al. 2009, Saavedra et al. 2015, Leprince et al. 2014, Meijer et al. 1999). In this study, we performed a genome wide analysis of the FYVE and PHOX gene families in *P. patens*. Gene structure and protein domain organization of these proteins is well conserved in comparison to higher plants. We performed a phylogenetic analysis and described these proteins in terms of their different domain composition and their potential binding site to PtdIns3*P*. Expression analyses based on RNAseq data together with the identification of cis-acting regulatory elements in promoter regions suggest the importance of these proteins particularly regulating developmental processes in *P. patens*.

## MATERIALS AND METHODS

### Identification of *FYVE* and *PHOX* genes

Sequences containing FYVE and PHOX domains from 11 plant species (Table S1) namely, *Chlamydomonas reinhardtii v5.5, Volvox carterii v2.1, Marchantia polymorpha v3.1, Physcomitrella patens v3.3, Sphagnum fallax v0.5, Selaginella moellendorffii v1.0, Arabidopsis thaliana, Glycine Max Wm82.a2.v1, Vitis Vinifera* and *Oryza Sativa* were retrieved using different approaches in Phytozome v12.1 (https://phytozome.jgi.doe.gov): a BLASTN search against the FYVE or PHOX genes previously described in *A. thaliana* and *O. sativa* [8-10], a BLASTP search using the FYVE domain amino acid sequence from the mouse hepatocyte growth factor-regulated tyrosine kinase substrate, Hrs [37], and through the BioMart data query tool (https://phytozome.jgi.doe.gov/biomart/martview) using the FYVE and PHOX domain PFAM accession terms PF01363 for FYVE and PF00787 for PHOX, respectively. *FYVE* and *PHOX* genes from *Klebsormidium nitens v1.1* was retrieved from the *Klebsormidium nitens* project (http://www.plantmorphogenesis.bio.titech.ac.jp/~algae_genome_project/klebsormidium/klebsormidium_blast.html). The presence of a FYVE or PHOX domain was further confirmed at the NCBI CCD search domain (https://www.ncbi.nlm.nih.gov/Structure/cdd/wrpsb.cgi), PFAM (http://pfam.xfam.org) and SMART (http://smart.embl-heidelberg.de) databases.

### Sequence alignment and phylogenetic analysis

Evolutionary analyses were conducted in MEGA7.0.26 (https://www.megasoftware.net). Multiple sequence alignment was inferred using MUSCLE. The evolutionary history was inferred using the Neighbor-Joining method and the bootstrap consensus tree was inferred from 1000 replicates. The evolutionary distances were computed using the Poisson correction method and are in the units of the number of amino acid substitutions per site.

### Chromosomal localization

The genomic localization of the FYVE and PHOX related genes through the 27 chromosomes of *Physcomitrella patens* was performed by KaryoPloteR R Bioconductor package (Gel and Serra 2017) using plotKaryotype and kpPlotMarkers functions. The custom genome parameter was created with toGRanges function from a BED file with the chromosome sizes.

### Promoter analysis

Promoter cis-acting regulatory elements within 1.5 kb of the upstream sequence from the ATG initial codon of each *P. patens FYVE* and *PHOX* gene were analyzed with PlantCARE (http://bioinformatics.psb.ugent.be/webtools/plantcare/html/). Analysis of transcription factor binding sites (TFBSs) of the 2.5 kb upstream sequence region of the initial codon was also performed using the Plant Promoter Analysis Navigator (PlantPAN2.0) (http://plantpan2.itps.ncku.edu.tw/), and with Homer v4.977 (http://homer.ucsd.edu/homer/motif/).

### Transcript data processing analysis

Raw sequencing data from *Physcomitrella patens* was survey in the Sequence Read Archive (SRA; https://www.ncbi.nlm.nih.gov/sra) and Gene Expression Omnibus (GEO; https://www.ncbi.nlm.nih.gov/geo/) databases from NCBI. Under transcriptomic data criteria, seven BioProjects were identified, five single-end (PRJNA294412, PRJNA153001, PRJNA149079, PRJNA265205) and two pair-end studies (PRJNA266515, PRJNA153109) (Table S10). The fastq files were download using fastq-dump command from NCBI C^++^ SRA Toolkit software version 2.10.0 (https://trace.ncbi.nlm.nih.gov/Traces/sra/sra.cgi?cmd=show&f=software&m=software&s=software) and quality control (QC) of raw reads was assessed with FastQC (https://www.bioinformatics.babraham.ac.uk/projects/fastqc/); trimming and adapter removal was performed using Trimmomatic (Bolger, Lohse, and Usadel 2014). Resulting reads were aligned to *Physcomitrella patens* (*P. patens* 318 v3.3) reference genome (https://phytozome.jgi.doe.gov/pz/portal.html#!info?alias=Org_Ppatens) using Kallisto (Bray et al. 2016). Mean quality scores report and summary statistics of read length, number of reads and read mapping are provided in Figure S3. Gene expression profiling was performed using edgeR (Robinson, McCarthy, and Smyth 2010) and limma (Ritchie et al. 2015) Bioconductor packages. The counts matrix obtained from Kallisto was read in edgeR, and the log_2_ transcripts per million (logTPM) was obtained and normalized using the trimmed mean of M values (TMM) method (Robinson, McCarthy, and Smyth 2010). Data were then filtered considering a fold change a FC > |1.5|.

### Network interaction of promoters

A co-expression network was built using the relationship of the gene vs known motif analysis (Homer v4.977; http://homer.ucsd.edu/homer/motif/) for PpFYVE and PpPHOX (Table S7 and Table S8). The gene and motif interaction matrixes were mapped employing Cytoscape 3.6.1 software (Shannon et al. 2003). Node table was generated using Molecular Complex Detection (MCODE) App (Bader and Hogue 2003) and nodes with more than four connections were depicted (Table S9).

## RESULTS

### Structural annotation of *PpFYVE* and *PpPHOX* genes

A search for genes coding for FYVE and PHOX domain proteins was performed in different photosynthetic organisms (*Chlorophytes*, *Charophytes*, *Bryophytes*, *Lycophytes* and *Angiosperms*) using different approaches (Materials & Methods). As a result, 136 FYVE and 88 PHOX domain-containing proteins were used in this study (Table S1). In the case of *P. patens* 13 and 9 FYVE and PHOX domain-containing proteins respectively, were identified in a total of 35,307 predicted protein-coding genes from the *P. patens* genome (Figure 1). The alternative splice variants, which shared a common genome locus and coding sequences, were tightly related to each other (Table S2 and S3), and therefore we selected the longer of the sequence variants for further analyses. *PpFYVE* and *PpPHOX* genes are dispersed throughout the *P. patens* genome, being located on different chromosomes (Figure 1) and the protein members of each subfamily have a conserved modular structure compared to their orthologs from other plant species (Figure 2).

**Figure 1.**
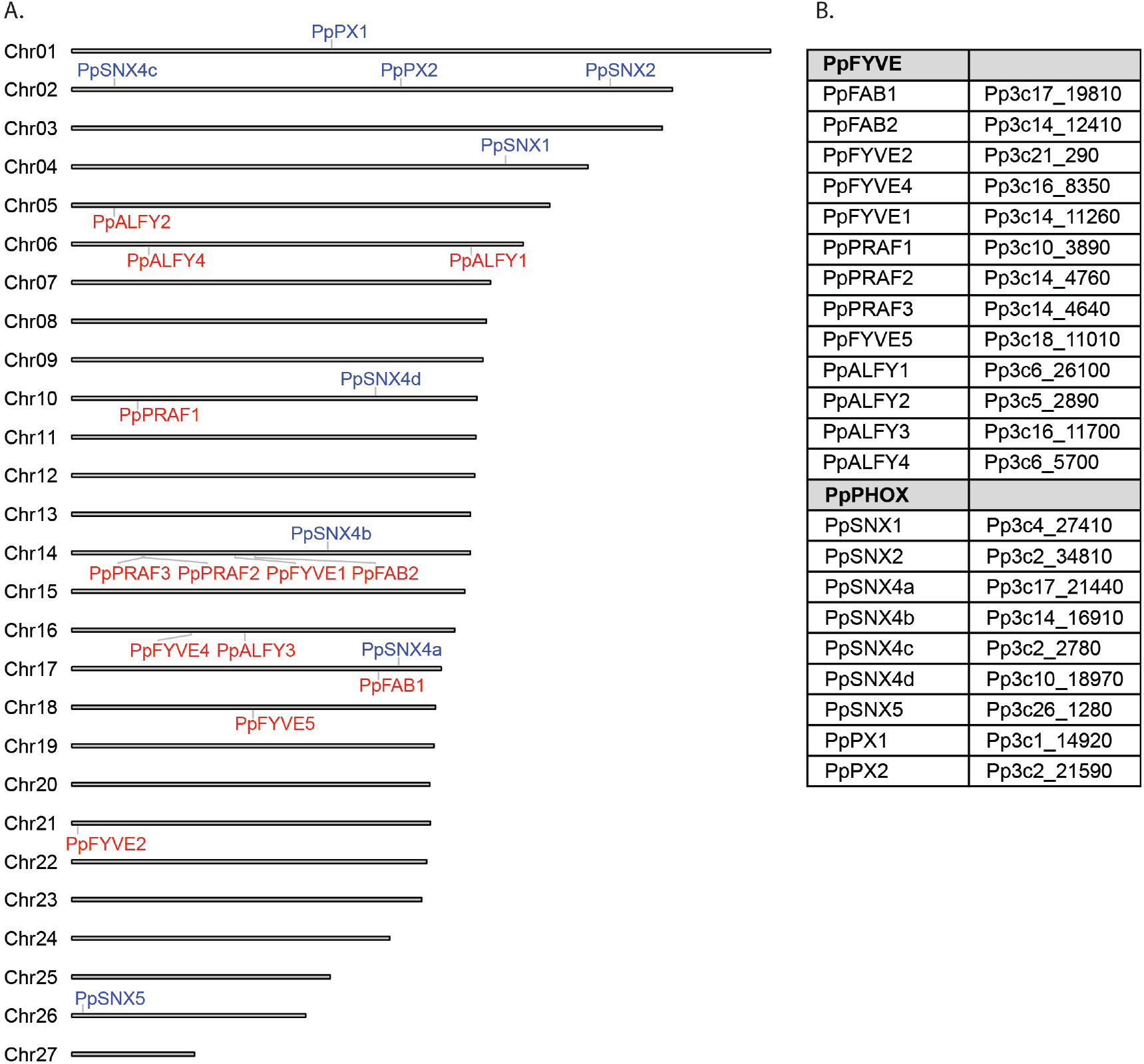
Chromosome localization and nomenclature of the *PpPFYVE* and *PpPHOX* genes in *P. patens*.

**Figure 2.**
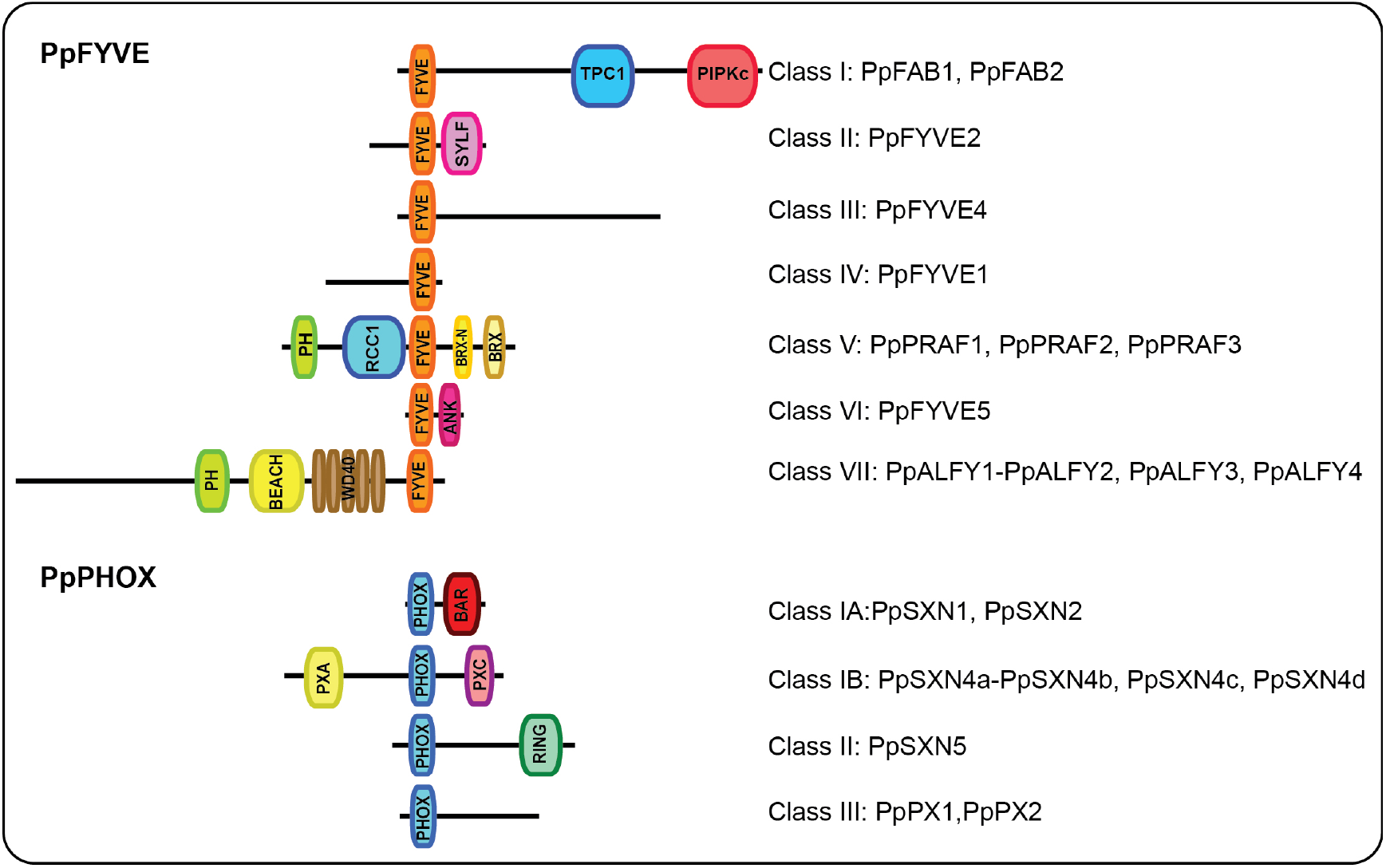
The FYVE and PHOX domain protein families in *P. patens*. Schematic diagram of the domain structures of the known plant FYVE and PHOX domain containing plant proteins. The nomenclature for the different proteins is shown in the right.

### *P. patens* FYVE proteins exhibit a similar modular structure in comparison to FYVE proteins from higher plants, but together with *Charophytes* exhibit an additional group

A phylogenetic analysis of PpFYVE proteins from the different organisms, as mentioned above resulted into 6 subfamilies previously described (Class I-VI) and an additional group (Class VII) with members only present in *Charophytes* and mosses (Figure 3). *Chlorophytes* exhibit very few FYVE proteins, one protein for Class II in *Chlamydomonas reinhardtii* and one protein for each Class II and I in *Volvox carterii*. Interestingly, *Klebsormidium nitens*, a filamentous terrestrial algae belonging to the *Charophytes*, which are widely accepted as the ancestors of current terrestrial plants exhibit one member for each of the seven classes of FYVE proteins identified in mosses. As soon as plants evolved to more complex forms the number of proteins for the different classes of FYVE proteins become in general larger, best exemplified by Class V, whereas the FYVE domain present in members of Class VII is lost.

**Figure 3.**
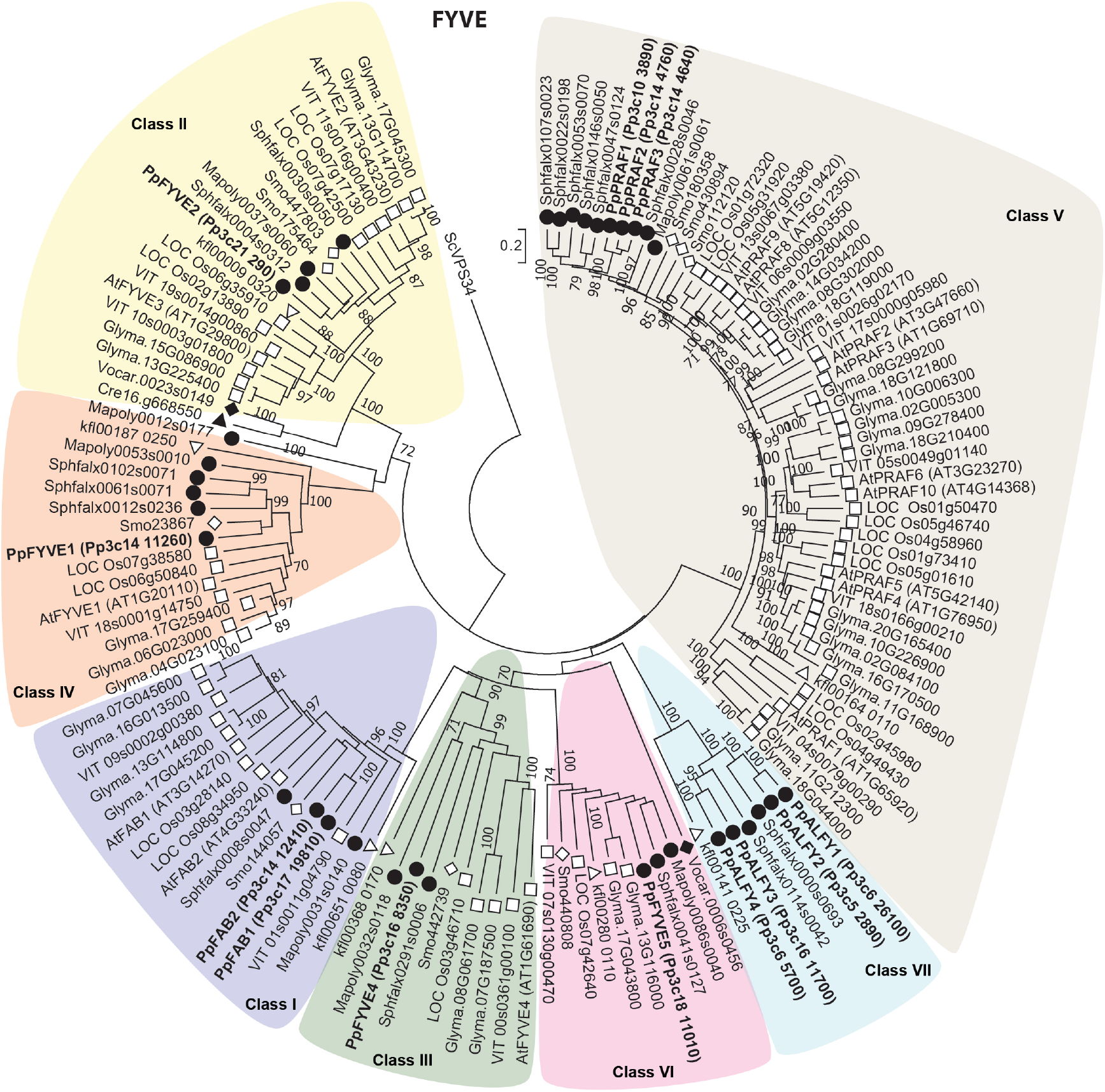
Phylogenetic analysis of FYVE proteins (classes I-VII) from *Chlorophytes (Chlamydomonas reinhardtii and Volvox carterii)*, *Charophytes (Klebsormidium nitens)*, *Bryophytes (Physcomitrella patens, Sphagnum fallax*, *Marchantia polymorpha)*, *Lycophytes* (*Selaginella moellendorffii*) and *Angiosperms (Arabidopsis thaliana, Glycine Max, Vitis vinifera and Oryza sativa)*.

Class I FYVE is represented by FAB kinases and has 2 members in *P. patens*, PpFAB1 and PpFAB2. FAB enzymes in plants, known as Fab1p in yeast and PIKfyve in mammals, are phosphatidylinositol 3-phosphate 5-kinases that synthesize PtdIns(3,5)*P*_2_ from PtdIns3*P*. Both PpFABs cluster together and share 81% similarity. They harbor three conserved domains: the N-terminal FYVE domain necessary for binding to PtdIns3*P*-containing membranes, a Cpn60_TCP1 (HSP chaperonin_T complex1) homology domain and a C-terminal kinase domain (Mueller-Roeber and Pical 2002), in accordance with the description of FABs (Figure 2). FAB proteins in plants have additional members lacking a FYVE domain. The single FAB protein present in *C. reinhardtii* lacks a FYVE domain, whereas *P. patens* and *A. thaliana* have one (Pp3c17_4720) and two members AtFAB1C (At1g71010) and AtFAB1D (At1g34260), respectively. The class II FYVE is represented by a single protein in *P. patens*, PpFYVE2 and consists of two, three and four members in *A. thaliana*, *V. vinifera* and *O. sativa*, respectively. Members of this family contain a FYVE domain and a C-terminal SYLF domain (also called DUF500). The SYLF domain is named “SYLF” on the basis of its representative members (SH3YL1, Ysc84p/Lsb4p, Lsb3p, and plant FYVE protein), and comprises a segment of ≈220 residues shown to be a lipid-binding module. Both the FYVE and the SYLF domain of AtFYVE2 also referred as CFS1 (CELL DEATH RELATED ENDOSOMAL FYVE/SYLF PROTEIN 1), were shown to bind to PtdIns3*P* (Sutipatanasomboon et al. 2017). A subgroup of proteins from class II including PpFYVE2 and AtFYVE2, exhibit a P(S/T)XP motif which is known from yeast and mammals to mediate binding to the Vps23/TSG101 subunit of the endosomal sorting complex required for transport (ESCRT)-I. AtFYVE2 is the only characterized protein from this class, and is involved in fusion events between autophagosomes and ESCRT-positive late endosomes (LEs), affecting autophagosome degradation and protein homeostasis (Sutipatanasomboon et al. 2017). The class III FYVE is represented by a single member in all species studied, except for *G. max* which has two members. *P. patens* PpFYVE4, contains only a FYVE domain close to the N-terminus and up to date no functional role has been reported for these proteins. Class IV together with Class I are the best characterized FYVE proteins in *A. thaliana*. Class IV FYVE harbors one member in *P. patens* (PpFYVE1) with only a FYVE domain localized close to the C-terminus. *FYVE1* has no homologs in animal cells, and in *A. thaliana* the homolog gene product named FREE1/FYVE1 binds to PtdIns3*P* through its FYVE domain and is involved in the degradation of ubiquitin-dependent membrane proteins, vacuolar transport, autophagy and vacuole biogenesis, showing FYVE1 as a central component of the traffic machinery, as stated previously (Gao et al. 2014, Barberon et al. 2014).

The largest subgroup of proteins with a FYVE domain in higher plants is Class V. It is absent in *Chlorophytes*, has one member in *K. nitens* and *M. polymorpha*, 3 members in *P. patens* (PpPRAF1-3) reaching to 9 members in *A. thaliana* and *O. sativa*. The three proteins present in *P. patens* have closer similarity to AtPRAF8 and AtPRAF9 (59-60% identity). Proteins from this group are characterized by two phosphoinositide binding domains, the PH (plekstrin homology) and FYVE domains, several repetitions of the RCC1 (regulator of chromosome condensation 1) motif, and a BRX (Brevis Radix) domain. PH domains bind various PPIs, but preferentially to PtdIns(4,5)*P*_2_, which was confirmed *in vitro* for the recombinant GST-PH_AtPRAF4_ (Jensen et al. 2001). Interestingly, all FYVE domains of PRAF proteins analyzed exhibit two important substitutions in the basic (R/K)_1_(R/K)HHCR_6_ motif, His4 and Arg6 which are components of the binding pocket that determines the specificity for binding of PtdIns3*P* (Gaullier et al. 2000) are substituted by asparagine and tyrosine (Supplemental figure 1). The RCC1 motif is present in several proteins in *P. patens* and in other organisms, it was reported as a versatile domain which may perform different functions, including guanine nucleotide exchange on small GTP-binding proteins, enzyme inhibition, or interaction with proteins and lipids (Hadjebi et al. 2008). A guanine nucleotide exchange activity has been demonstrated for AtPRAF1 (Jensen et al. 2001). The BRX motif is a protein-protein interaction domain, and it is present as tandem repeats in AtBRX, a transcription factor that regulates cell growth (Beuchat et al. 2010, Mouchel, Briggs, and Hardtke 2004). A FYVE domain followed by an ankyrin domain characterizes the modular structure of class VI FYVE proteins. Interestingly, this class is absent in *Chlorophytes* and in *A. thaliana* although is present in the other organisms analyzed in this study. The ankyrin domain mediates protein-protein interactions, but no functional role of this subgroup has been addressed yet.

Finally, members of the last class of FYVE-domain proteins, class VII from the organisms analyzed were only present in *K. nitens*, and in the mosses *P. patens* (PpALFY1-4) and *S. fallax*. These proteins exhibit a similar domain structure to the mammalian protein ALFY (<u>a</u>utophagy-linked <u>FY</u>VE protein), which is required for degradation of protein aggregates by autophagy (aggrephagy) (Simonsen et al. 2004). Proteins belonging to this subgroup exhibit in addition to the FYVE domain located close to the C-terminus and a PH domain followed by a BEACH domain and a tandem repeat of WD domains. Interestingly, proteins lacking the FYVE domain but with a similar modular structure are present in *M. polymorfa, S. moellendorffii* and all the angiosperms analyzed in this study, suggesting that this domain was lost during higher plants evolution. The BEACH domain consists of ∼280 amino acids and is found in eukaryotic proteins involved in vesicle trafficking, yet the function of this domain is unknown. The WxxD, RRHHCR and RVC motifs are less conserved in PpALFYs, any of the four proteins conserved the WxxD motif, and PpALFY 1-2 are the two members with a conserved RRHHCR and RVC motifs (Supplemental Fig.1).

### *P. patens* PHOX proteins are classified into 5 groups, share a similar modular structure in comparison with PHOX proteins from higher plants but lack PLDζ with this domain

The phylogenetic analysis of PHOX proteins from the *Chlorophytes*, *Charophytes*, *Bryophytes*, *Lycophytes* and *Angiosperms* analyzed, grouped these proteins into five different groups named Class I-V (Figure 2 and Figure 4). Members from *P. patens* clustered with the subfamilies previously described for *A. thaliana* with the exception of class V, which are phospholipases D type Zeta (van Leeuwen et al. 2004).

**Figure 4.**
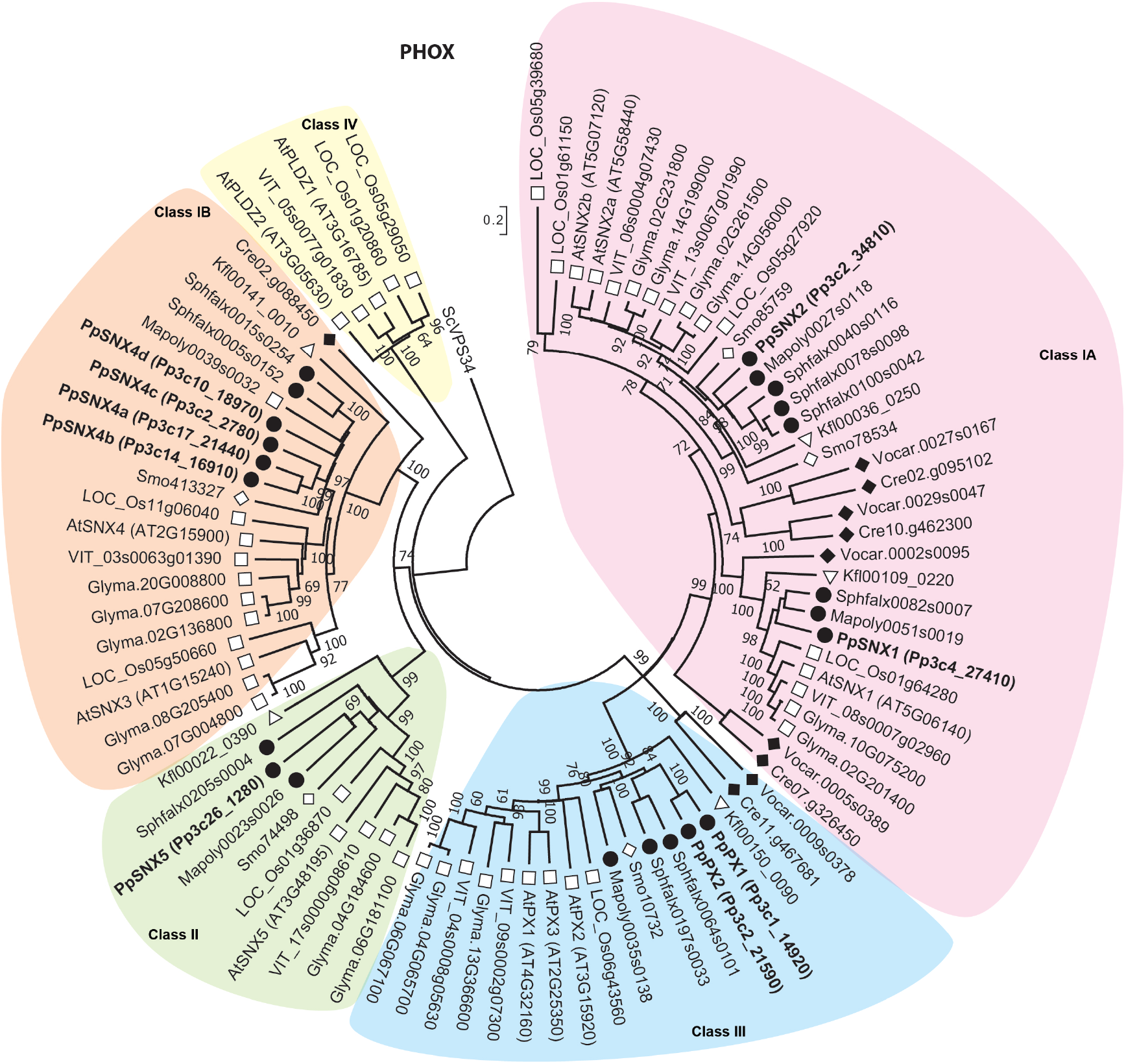
Phylogenetic analysis of PHOX proteins (classes I-IV) from *Chlorophytes (Chlamydomonas reinhardtii and Volvox carterii)*, *Charophytes (Klebsormidium nitens)*, *Bryophytes (Physcomitrella patens, Sphagnum fallax*, *Marchantia polymorpha)*, *Lycophytes* (*Selaginella moellendorffii*) and *Angiosperms (Arabidopsis thaliana, Glycine Max, Vitis vinifera and Oryza sativa)*.

The proteins belonging to classes I and IV are represented by the SNX (Sorting Nexin) protein family. Several SNXs are components of the retromer complex, which is involved in the recycling/retrograde transport of VSRs (Vacuolar Sorting Receptors) and PM proteins from endosomes to the trans Golgi network (TGN) (Heucken and Ivanov 2018). Class I is composed of two members: PpSXN1 which shares 49% identity with AtSNX1 (AT5G06140), and PpSNX2 which clusters with AtSNX2a (AT5G58440) and AtSNX2b (AT5G07120) and shares with them 56.89% and 55.40 % identity, respectively. In addition to the PHOX domain, class I proteins exhibit a BAR (Bin-Amphiphysin-Rvs) domain at their C-terminal region which enables dimerization and binding to curved membranes. The class II is represented by PpPX1 and PpPX2, and exhibit only a PHOX domain positioned close to the N-terminus. *A. thaliana* has three members within this group, AtPX1/EREL1 (AT4G32160), AtPX2/EREX (AT3G15920) and AtPX3/EREL2 (AT2G25350). The AtEREX (Endosomal Rab Effector with PX-domain) protein has been suggested as a genuine effector of canonical RAB5s mediating vacuolar trafficking (Sakurai et al. 2016). Members of class III are single proteins in all species analyzed with exception of *G. max*, and are represented by PpSXN5 in *P. patens*. These proteins exhibit a PHOX and a C-terminal RING9 domain, but none physiological role has been assigned to this group yet. Proteins from class IV in *P. patens* (PpSNX4a-d), contain a PXA (PHOX-associated) domain, the PHOX domain and a conserved C-terminal domain, referred to as PXC domain. *A. thaliana* and *P. patens* PHOX domains from this group have homology to human SNX19 PHOX domains, but the function of these proteins in plants remains unknown. Class V contains members of phospholipase D type Zeta (PLDζ: AT3G16785 and AT3G05630). By using the BioMart data query tool with the PFAM accession term for PHOX domain proteins in *P. patens* none PLD sequences were obtained. Therefore, protein sequences with a high percentage of identity to AtPLDζ were retrieved, resulting two sequences Pp3c13_3440V3.1 and Pp3c12_3620V3.1, which were further analyzed. None of them scored a PHOX domain associated with a significant *E*-value as noticed for PHOX domains from HsPLD1 and HsPLD2 (Supplemental Table 4). Thus, *P. patens* PLD sequences were excluded from the analysis.

Additionaly, an *in silico* analysis was performed to search if PHOX domains from *P. patens* exhibit the amino acids required for the binding to PtdIns3*P.* Bravo et al. (Bravo et al. 2001) have previously described the crystal structure of the p40^phox^ PHOX domain bound to PtdIns3*P*, denoting the four key residues within this domain, Arg58, Tyr/Phe59, Lys92 and Arg105 (numbers corresponding to p40^phox^ PHOX as reference) that determine specificity and are required for binding to PtdIns3*P.* Analysis of these sites in the PpPHOX family showed that all members of Class I and II harbor these four key residues (Supplemental Figure 2), suggesting that these domains could bind PtdIns3*P in vivo*. The exception to this, was exhibited by proteins of Classes III and IV, PpSNX4a-d and PpSNX5, where Lys92 is substituted by arginine, but still conserved the positive charge.

### *Cis-*acting regulatory elements in promoter regions

Analysis of *cis*-regulatory elements from the *PpFYVE* and *PpPHOX* promoter regions was performed using the PlantCARE (Table S5), PlantPAN database (Table S6) and Homer software (Tables S7 and S8). *PpFYVE* and *PpPHOX* promoters contain in addition to the core *cis*-elements, including the TATA box and CAAT box motifs presented in all promoter regions, several regulatory motifs identified and associated with light regulation (BOX I, BOX 4, ACE, MRE, LAMP), low temperature and drought responses (LTR, MBS), defense and stress responses (e.g. TC-rich repeats), hormonal regulation such as salicylic acid (e.g TCA-element), methyl jasmonate (e.g CGTCA-motif, TGACG-motif), auxin (AuxRR-core, TGA-element), abscisic acid (ABRE), and regulatory motifs related to developmental processes/ cell differentiation (HD-Zip 1, HD-Zip 2). A transcription factor (TF) survey of binding sites was performed to FYVE and PHOX genes using the Homer platform, and elements with a score higher than 10 were used for further analysis. Interestingly, the classes of TFs binding sites identified in the promoter region of various members from *PpFYVE* and *PpPHOX* genes were those associated mainly with plant growth and developmental processes and with stress responses, such as ARID, ABI3VP1, C2C2 dof, C2H2, NAC, G2 like, Homeobox (HD-ZIP), Myb (Figure 5 and Figure 6). Several TFs binding sites were present in promoters from a subset of both *PpFYVE* and *PpHOX* genes including ATHMGB15 (ARID, AT1G04880), AT2G40260 (G2like), ATHB13, ATBH51, ATBH53 and HGDI (Homebox), AT5G60130 (ABI3VP1), IDD5(C2H2) and At4g38000 (C2C2dof) (Table S9).

**Figure 5.**
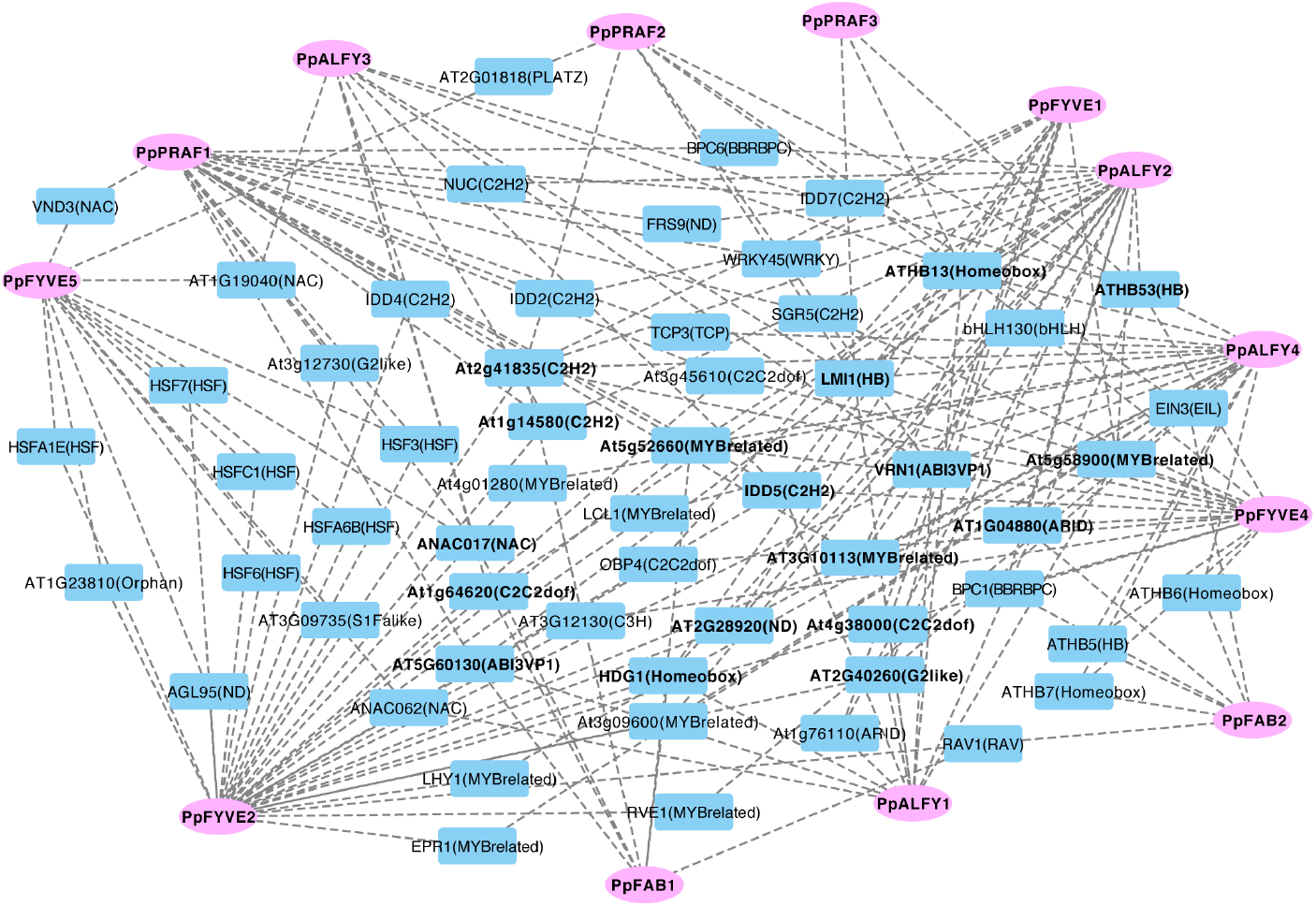
Gene-motif interaction network of *PpFYVE* genes. Motifs were filtered by a motifScore > 10 and a degree > 4. Nodes are represented as *PpFYVE* transcripts (in pink) and motifs (in blue), edges show their interaction (gray dash line).

**Figure 6.**
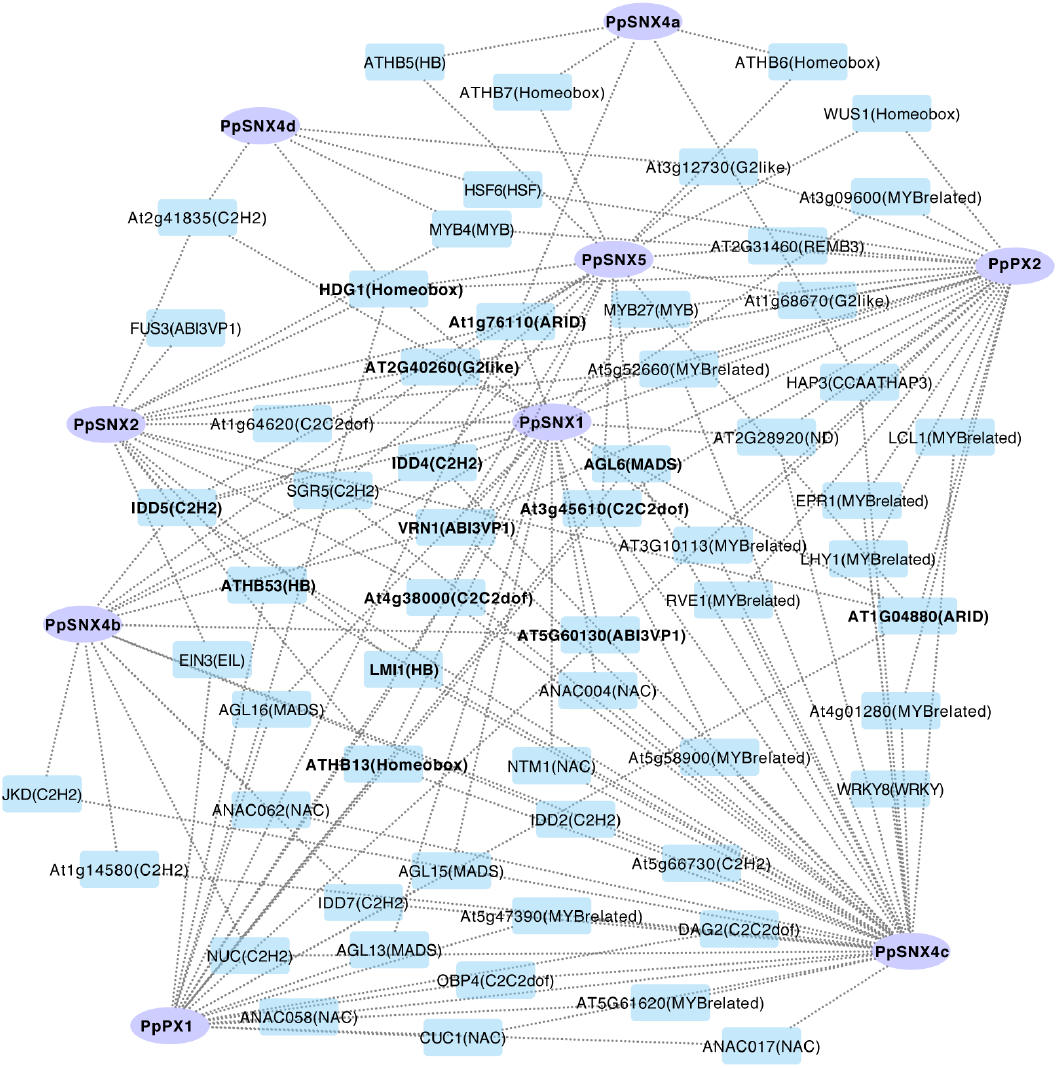
Gene-motif interaction network of *PpPHOX* genes. Motifs were filtered by a motifScore > 10 and a degree > 4. Nodes are represented as *PpPHOX* transcripts (in purple) and motifs (in light blue), edges show their interaction (gray dash line).

### Expression analysis of *P. patens FYVE* and *PHOX* genes

#### Gene expression during development

The transcriptional signature of *PpFYVE* and *PpPHOX* was studied among different developmental stages of *P. patens* life cycle and under abiotic stress conditions (Table S8 see Materials and Methods). As it is briefly described herein, *P. patens* life cycle is dominated by a photoautotrophic haploid gametophytic generation, that supports an ephemeral heterotrophic diploid sporophyte (Cove et al. 2006). *P. patens* gametophyte consists of a juvenile filamentous protonemata and the leafy adult stage (gametophore or leafy shoot). Protonemata cells, chloronemata and caulonemata, develop by apical growth and cell division of apical and subapical cells. Chloronemata are the first cells to emerge from the spore or protoplast, whereas caulonemata develop by progressive differentiation of chloronemal apical cells in response to environmental signals such as light and auxin. Caulonema cells form side branch initial cells of which a few become buds (gametophore apical cells), which are meristematic and develop into an adult gametophore comprising a photosynthetic non-vascularized stem, leaves, the reproductive organs, supported by the development of rhizoids. Protonemata and rhizoids are cell types that exhibit polar growth.

During the different stages of moss development, several *PpFYVE* and *PpPHOX* were modulated revealing the importance of these proteins in regulating developmental processes (Figure 7). The data used for this study comprises different time points during protoplast regeneration (24 h, 48 h and 72 h) that were compared against 0 h. Within the first 24 h of culture a new cell wall is developed, new polar axes are re-established within 48 h when reprogramming into stem cells occurred, and after 72 h protoplasts divided asymmetrically leading to chloronemata that contained two to three cells (Xiao et al. 2012). During this period of time several *FYVE* genes were down-regulated, namely *PpFAB2*, *PpFYVE4*, *PpFYVE1*, *PpALFY1* and *PpPRAF3* (Figure 5A). Also, these genes were down-regulated in the comparison of chloronema vs protoplast in the expression data from the eFP Physcomitrella Browser (Ortiz-Ramírez et al. 2016). In juvenile protonemata *PpFAB1-2*, *PpPRAF2*, *PpALFY3* were up-regulated in the comparison caulonemata vs chloronemata, whereas *PpFYVE4* and *PpALFY3* along with *PpFYVE1* were up-regulated in the bud vs tip comparison. *PpPRAF3* was down-regulated at several stages of development (3, 14, 24 and 30 d), while *PpALFY3* was up-regulated at 24 days, time point when buds had formed from protonemal filaments and represents initiation of leafy shoots and at 30d (Xiao et al. 2011). *PpALFY1* was the only *FYVE* gene up-regulated when gametophore was compared to the whole plant, and both *PpALFY4* and *PpFYVE5* were up-regulated in the young sporophyte comparing to the whole plant. The expression patterns observed at 24 and 30 days compared to 3 days and chloronemata, also suggest that a group FYVE genes (*PpPRAF2, PpALFY3, PpFYVE4*) may acquire a role at latter stages of moss development, during the two dimensional (2D) to three dimensional (3D) growth transition in a moss.

**Figure 7.**
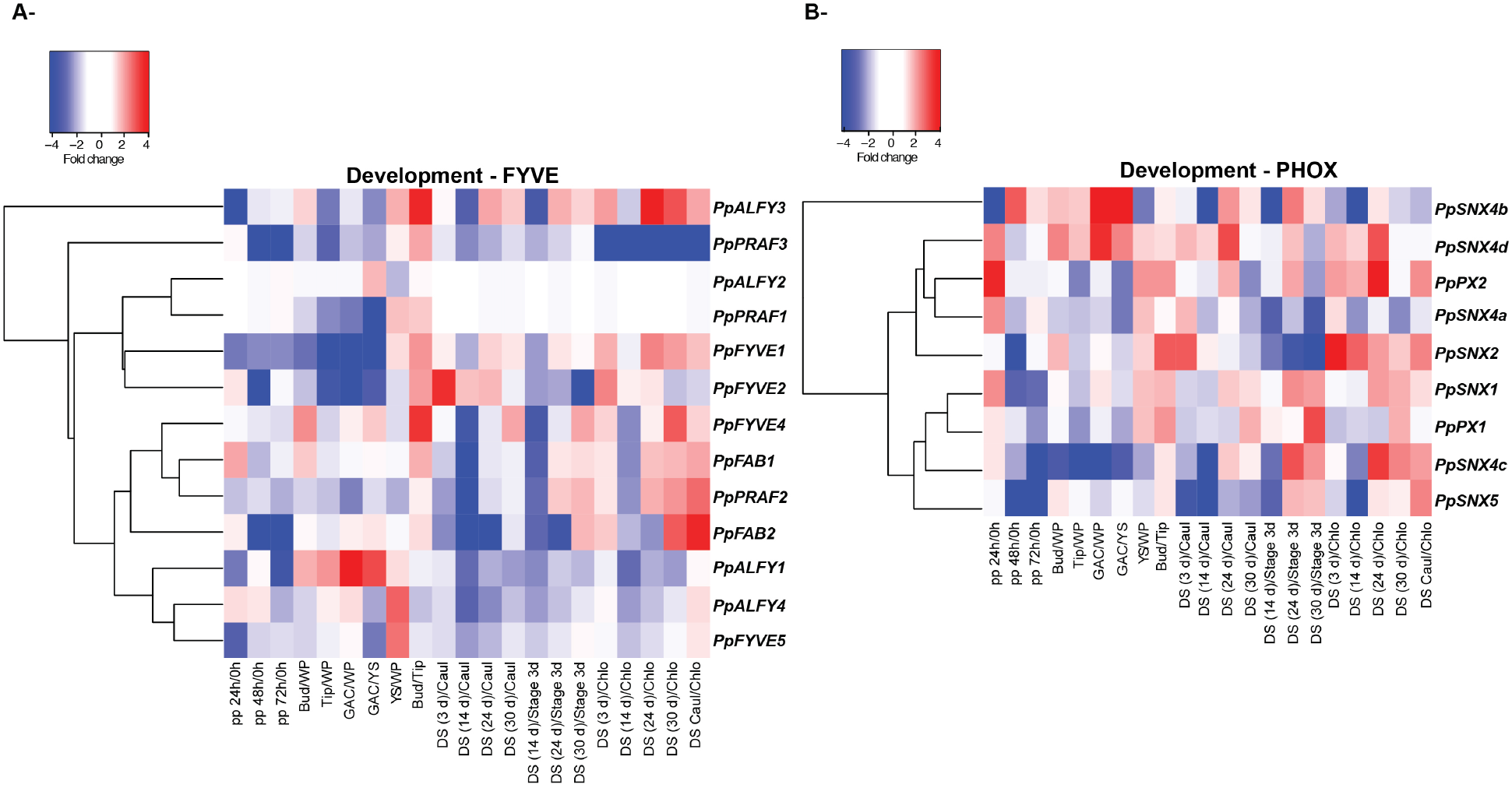
Heatmap analysis of the *PpFYVE* and *PpPHOX* genes during developmental stages of *P. patens* life cycle. Gene expression of *PpFYVE* and *PpPHOX* were assessed in three RNAseq studies (columns) and a hierarchical cluster analysis was applied at transcript level (rows). Studies from left to right: protoplast (PRJNA153001; Xiao, et al 2012), developmental tissues (PRJNA265205; Frank, et al 2015) and developmental stages (DS) (PRJNA149079; Xiao L., et al 2011). Comparisons are denoted by the character [/]. Key color represents the fold change values, up-regulated genes (red) and down-regulated (blue). Abbreviations: pp, protoplast; WP, whole plant; GAC, gametophore apical cell; YS, young sporophyte; DS, developmental stage; Caul, caulonema; Chlo, chloronema.

In the case of the expression of *PHOX* genes, *PpPX2* together with *PpSNX1*, *PpSNX4a* and *PpSNX4d* were up-regulated at 24 h vs 0 h of protoplast regeneration (Figure 5B). However, expression of *PpPX2, PpSNX1* together with *PpSNX5* and *PpSXN4c* decreased at latter time points of this process (72h compared to 0 h) when chloronema development was initiated. During protonemata development the syntaxins *PpSNX2*, *PpSNX5* and *PpPX2* were up-regulated in caulonemata compared to chloronemata. *PpSNX4d*, *PpSNX2* were up-regulated in the comparison of bud vs whole plant or tip. This result is reinforced by the fact that these genes are also up-regulated at 24 days (stage where buds had developed from protonemal filaments) compared to chloronemata. Several *PHOX* genes were down-regulated in the comparison chloronemata against caulonemata (development stage 14 days composed of protonemata where chloronemal cells were about twice as numerous as caulonemal cells), namely *PpSNX4b, PpSNX4c* and *PpSNX5*. Additionally, *PpSNX4c* presents a strong down-regulation in several cell types but it is up-regulated in the developmental stage of 24 days rich in caulonemata. *PpSNX4b* and *PpSNX4d* were up-regulated in the gametophore comparing to the whole plant and *PpPX2* was up-regulated in the young sporophyte comparing to the whole plant.

#### Gene expression upon abiotic stress

Expression signature under abiotic stress of *PpFYVE* and *PpPHOX* revealed that the majority of their transcripts were down-regulated upon the stresses imposed by ABA, cold, salt, light and drought treatments (Figure 8). As example, *PpFAB1*, *PpALFY1*-3*, PpPRAF2* and *PpFYVE5* were down-regulated in drought conditions (30 min) but up-regulated in the *anr (ABA NON-RESPONSIVE)* mutant (Figure 6A), which fail to express dehydration tolerance-associated genes in response to drought, ABA, or osmotic stress and do not acquire ABA-dependent desiccation tolerance (Stevenson et al. 2016). Additionally, *PpALFY1,2 and 4* genes were down-regulated upon salt treatment. While, *PpSXN4c* was the only *PHOX* gene up-regulated during ABA, cold and salt treatments after 30 min (Figure 8B).

**Figure 8.**
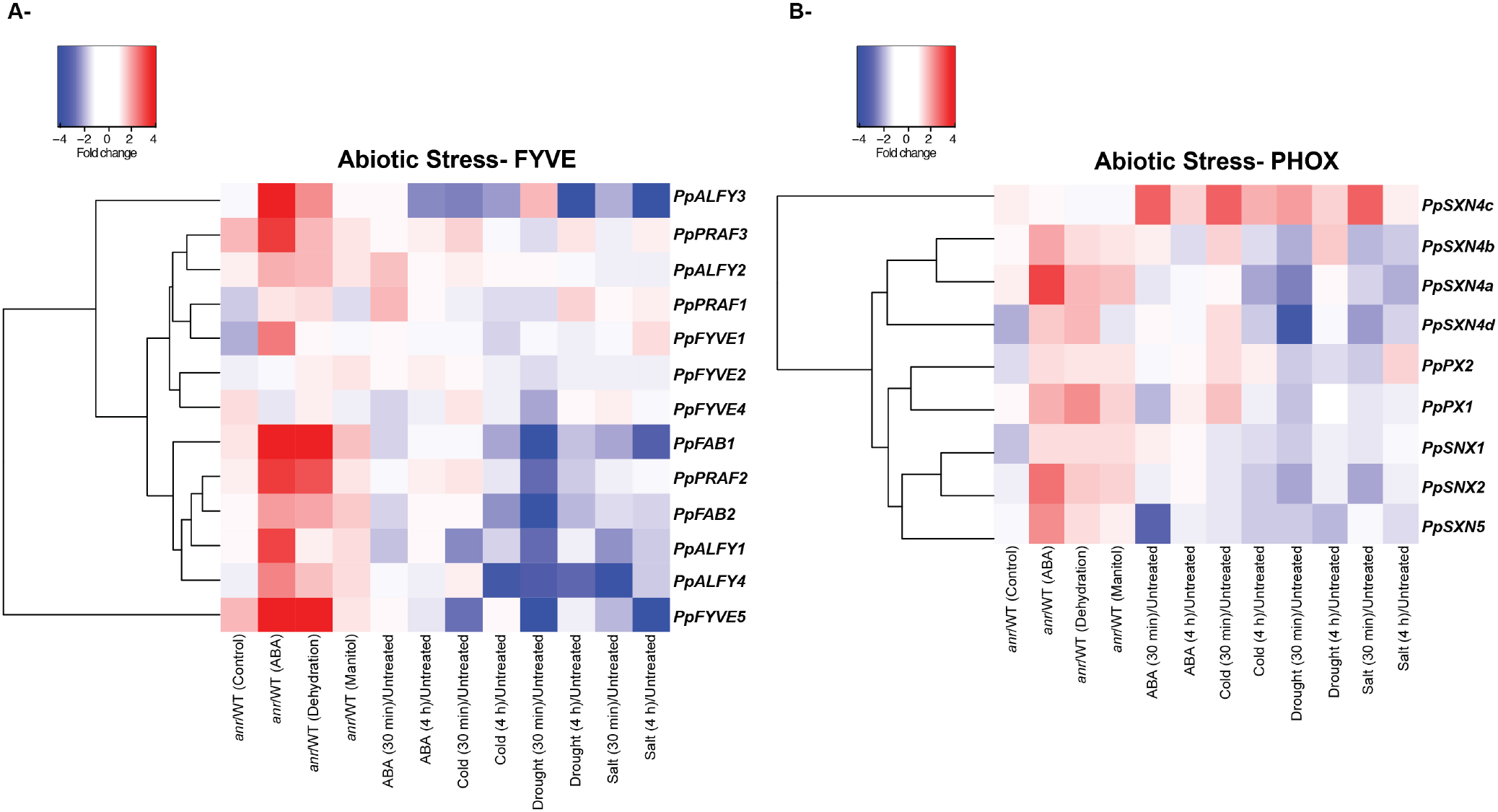
Heatmap analysis of the *PpFYVE* and *PpPHOX* genes under different abiotic stress treatments in *P. patens*. Gene expression of *PpFYVE* and *PpPHOX* were assessed in two RNAseq studies (columns) and a hierarchical cluster analysis was applied at transcript level (rows). Studies from left to right: *anr* mutant (PRJNA294412; Stevenson et al 2016), abiotic treatments through the time (PRJNA266515). Comparisons are denoted by the character [/]. Key color represents the fold change values, up-regulated genes (red) and down-regulated (blue). Abbreviations: *anr* (aba non-responsive), WT (Wild Type), and ABA (abscisic acid).

## DISCUSSION

PtdIns3*P* is synthesized from PI by type III PI3Ks [(phosphatidylinositol 3-kinase, also known as VPS34 (Vacuolar Protein Sorting 34)]; the only type of PI3K present in plant genomes. Either by silencing the *PI3K* gene, or by using the pharmacological type III PI3K inhibitors, it was revealed that PtdIns3*P* is involved in several biological processes such as the polar growth of root hairs (Lee, Bak, et al. 2008), pollen development (Lee, Kim, et al. 2008), auxin induced root gravitropism (Joo et al. 2005) and seed germination (Liu, Zhou, and Xing 2012). Under stress conditions, PtdIns3*P* is required for stomatal closure (Park et al. 2003). In response to salt stress, it is necessary for endocytosis of PM and intracellular ROS production within endosomes (Leshem, Seri, and Levine 2007), and regulates proline metabolism (Leprince et al. 2014). Furthermore, it has been shown to be essential in the early stages of mutualistic interactions of legumes with beneficial microorganisms (Peleg-Grossman, Volpin, and Levine 2007, Robert et al. 2018, Estrada-Navarrete et al. 2016), as well as in the pathogenic responses (Kale et al. 2010). However, the mechanistic insights underlying PtdIns3*P* mode of action, specifically downstream targets of PtdIns3*P*, still remains partially unexplored in plants. Moreover, PtdIns3*P* is the precursor of PtdIns(3,5)*P*_2_, which has also important roles in membrane trafficking (Hirano, Munnik, and Sato 2017), thus the importance to distinguish between the roles attributed to PtdIns3*P* or due to an inability of cells to produce PtdIns(3,5)*P*_2_ due to the lack of its precursor. Therefore, unraveling the functional role of PtdIns3*P* effectors such as FYVE and PHOX-domain proteins would provide advances in this area.

### Evolution of FYVE and PHOX domains in Viridiplantae

Previous studies have shown that FYVE and PHOX domains are distributed across various eukaryotic lineages, but with a distinct selectivity. While Fungi and Metazoa groups have a higher occurrence of the PHOX domain, Streptophyta of Viridiplantae, exhibit an opposite trend, with a higher number of FYVE domain proteins (Banerjee, Basu, and Sarkar 2010). Phylogenetic analysis performed in this study divided the FYVE gene family into 6 subfamilies previously described (Class I-VI), and an additional group (Class VII). Members of the *Chlorophytes* such as *C. reinhardtii* and *V. carterii* have only Class II and Class I/II FYVE proteins, respectively (Figure 3), whereas other members such as *Ostrococcus lucimarinus* and *Ostreococcus tauri* do not encode any FYVE proteins (Banerjee, Basu, and Sarkar 2010). Class I (FABs), are highly conserved proteins from yeast to mammals and plants, and members from this class conserved all the canonical ligand-binding site consensus sequence of RRHHCR (Supplemental Fig. 1). Interestingly, although *C. reinhardtii* lack FAB proteins with a FYVE domain, it harbors the capacity to synthetize PtdIns(3,5)*P*_2_ (Meijer et al. 1999), and in similar fashion, *in vitro* activity assays using heterologous recombinant AtFAB1C-GST, which lacks a FYVE domain shown its ability to synthetize PtdIns(3,5)*P*_2_ from PtdIns3*P* (Bak et al. 2013). This suggests that other amino acids residues different from those of the FYVE domain are important/required for substrate recognition and specificity for the catalytic activity of these proteins.

The class II of FYVE proteins is present in all species analyzed. The first arginine in the consensus sequence R_1_RHHCR has a substitution for glycine or serine (Supplemental Fig. 1), but in *A. thaliana* it maintains the ability to bind PtdIns3*P*, together with the SYLF domain exhibited by these proteins (Sutipatanasomboon et al. 2017). A subgroup of proteins from class II already present in *Chlorophytes* which includes PpFYVE2, harbor a P(S/T)XP motif, which mediate binding to the Vps23/TSG101 subunit of ESCRT-I. AtFYVE2, the *A. thaliana* ortholog of PpFYVE2 acts as an autophagy regulator and repressor of cell death, it binds to the ECL component of ESCRT-I LEs and is involved in autophagosome turnover (Sutipatanasomboon et al. 2017).

Through the analysis performed in *K. nitens*, our results indicate that the seven classes of FYVE domain proteins are present in *Charophytes* (Figure 3), the closest living relatives of land plants. The data suggest that different to Classes I and II, which evolved earlier, PtdIns3*P* signaling through Classes III-VII is involved in processes necessary for terrestrialization, ej. acquisition of adaptive mechanisms to cope to abiotic stresses and/or the evolution to more complex body sizes through cellular differentiation. A suitable example in this regard, is the Class V (PRAFs), which are plant-specific proteins, characterized apart from the FYVE domain, by a PH, RCC1 and BRX domains. The single functional report of these proteins showed its requirement for normal root and nodule development in *M. truncatula* (Hopkins et al. 2014). Interestingly, the BRX domain of AtPRAF4 (At1g76950) was shown to interact with BRX proteins, which are only found in multicellular plants, and control the extent of cell proliferation and elongation in the growth zone of the root tip (Briggs, Mouchel, and Hardtke 2006). PRAF proteins exhibit another deviation of the RRHH_4_CR_6_ canonical motif, where asparagine and tyrosine substitute His4 and Arg6 (Supplemental Fig. 1). Although a recombinant 6XHis-FYVE_AtPRAF4_ exhibit binding towards PtdIns3*P*, a double mutant GST-FYVE_AtPRAF4-N658H,Y660R_ has an increased affinity to PtdIns3*P*, highlighting the importance of this residues for the specific binding (Gaullier et al. 2000, Jensen et al. 2001). Yet, the role of FYVE domain in these proteins is unknown.

Class VII proteins were only found in *K. nitens* and in *P. patens*. Although the proteins with a modular structure of PH, BEACH and WD40 domains are conserved through land plant evolution, the FYVE domain is absent in these proteins in the liverwort *M. polymorpha* and in tracheophytes, suggesting that its function was not further essential. Interestingly, the FYVE domain of PpALFYs exhibits a low degree of conservation in comparison to the canonical FYVE sequence (Supplemental Fig. 1).

Phylogenetic analysis of PHOX proteins clustered these proteins into five different groups, Class I-V. Different to what was observed for FYVE proteins, PHOX-domain proteins for classes I, II and IV, with exception for class III and V (PLDζ), are already present in *Chlorophytes*, suggesting that these proteins are involved in conserved processes of eukaryotic cells through evolution, reported as both cargo adaptors in cell trafficking and scaffolds regulating signal transduction.

The subfamily of Class I proteins possessing a C-terminal BAR domain are components of the retromer, a key protein complex involved in cargo recycling and retrograde transport. Different to what observed in mammalian cells, SNXs are dispensable for membrane binding and function of the retromer complex in *A. thaliana* (Pourcher et al. 2010). Components of the core subunit of the retromer (Vps35p, Vps29p and Vps26p) can work with or independently of SNXs in the trafficking of seed storage proteins, suggesting distinct functions for subcomplexes of the plant retromer, or that these proteins perform additional functions independently of the it (Heucken and Ivanov 2018, Pourcher et al. 2010). A functional role for proteins from class II was recently described in *A. thaliana*, with AtEREX as a genuine effector of canonical RAB5s and the two homologs (AtEREL1 and AtEREL2) play partially redundant functions in the transport of seed storage proteins (Sakurai et al. 2016). The role of proteins from class III and IV is unknown. Interestingly, proteins from class III have in addition to the PHOX domain a C-terminal RING9-type Zn-finger domain, but this modular arrangement has not been found in other organisms, suggesting plant-specific functions.

All PHOX domains from *A. thaliana* and *P. patens* studied exhibit the four residues R_58_Y/F_59_K_92_R_105_ (with p40^phox^ as reference (Bravo et al. 2001)), critical for PtdIns3*P* recognition conserved, with exception of members form Classes III and IV, where Lys92 is substituted by another positively charged amino acid, arginine. The PHOX domain specificity towards PPIs has been tested for AtSNX2b, AtEREX (At3g15920) and AtSNX1. AtSNX2b and AtEREX binds to PtdIns3P in vitro, and this association with the former is required for the localization of AtSNX2b to punctate structures in vivo, identified as the *trans*-Golgi network, prevacuolar compartment and endosomes (Phan, Kim, and Bassham 2008). In the case of AtSNX1, although it binds to PtdIns3P, it prefers PtdIns(3,5)P_2_ *in planta* (Pourcher et al. 2010, Hirano, Munnik, and Sato 2015).

### *PpFYVE* and *PpPHOX* promoters contain several TFs binding sites involved in development of pollen and root growth

Analysis of putative TFs binding sites in *PpFYVE* and *PpPHOX* promoters showed groups of genes with same TF binding sites with interesting functions in growth and development. For example, *PpSNX1*, *PpSNX2*, *PpSNX4c*, *PpPX1* have all sites for homeodomain leucine zipper class I *HB13*, *HB53* and *HB51*. In *A. thaliana HB13* is crucial for pollen germination (Capella, Ribone, and Chan 2015), *HB53* suggested to have a regulatory role in auxin/cytokinin signaling during root development (Son et al. 2005), and *HB51* (*LMI1*) is a meristem identity regulator (Saddic et al. 2006). Interestingly, these three TF binding sites were also observed in promoters of several PpFYVE as *PpPRAF2-3*, *PpALFY1-4* and *PpFYVE4*. *PpSNX1*, *PpSNX4c*, *PpPX1* cluster together in the heatmap of developmental stages in *P. patens* inviting to explore (Figure 7B) the role of these genes in protonemata and rhizoids apical growth and during the development of meristematic buds. Another TF binding site observed in promoters from *FYVE* and *PHOX* genes is *HDG1*, which belongs to the HD-ZIP IV family and promote cell differentiation (Horstman et al. 2015), in addition to BLUEJAY and RAVEN which are part of the network regulated by BLJUEJAY, JACKDAW, SACRECROW and SHORT-ROOT to regulate root tissue patterning through cell lineage specification and asymmetric cell division (Moreno-Risueno et al. 2015).

### *PpFYVEs* are specifically expressed in different developmental stages of *P. patens*, and several members are up-regulated in caulonemata and gametophore apical cells

*FYVE* and *PHOX* genes showed different expression patterns across the developmental stages of *P. patens*, but until now none of these genes have been characterized in *P. patens* with exception of *PpFABs*, where silencing these genes impaired protonema polarized growth (van Gisbergen et al. 2012). Aside from Class I (*FABs*), *PpFYVE* classes with additional members as Classes V and VII did not exhibit a subclass-specific expression pattern, suggesting gene-specific function or localization regardless the phylogenetic class. *PpFABs* are up-regulated in caulonemal and bud cells, whose differentiation and formation, respectively, is stimulated by auxin (Thelander, Landberg, and Sundberg 2018). Interestingly, pleiotropic auxin-signaling phenotypes are observed in *Atfab1a/b* mutants due to a mislocalization of auxin transporters, as a result of the impairment of the recycling process of these proteins between the PM and endosomal compartments (Hirano et al. 2011). Furthermore, synthesis of PtdIns(3,5)*P*_2_, FABs synthesis product, on LEs is necessary as a scaffold for the recruitment of LE effector proteins for endosome maturation, giving the identity of LE necessary for the proper PIN trafficking (Hirano, Munnik, and Sato 2015). In close relation to this role for organelle maturation/acidification, synthesis of PI(3,5)*P*_2_ by FABs is necessary for vacuolar acidification in pollen tubes (Serrazina, Dias, and Malhó 2014), and in guard cells during stomatal closure induced by ABA in *A. thaliana* (Bak et al. 2013). The tonoplast proton transporter V-PPase was suggested as a putative effector in the control of vacuolar acidification since it binds PtdIns(3,5)*P*_2_ *in vitro* (Bak et al. 2013). Another potential effector of PtdIns(3,5)*P*_2_ is the V-ATPase, which is critical to ensure efficient phagosomal acidification in the defense against pathogenic microbes (Buckley et al. 2019). It would be of particular interest to evaluate if FABs have a role in the autophagic process in plant cells, in particular related to achieve/maintain the required acidic environment in autophagic vesicles and in the lytic vacuole for proper functioning.

*PpFYVE1*, *PpPRAF2* and *PpALFY3* are *PpFYVEs* that were also up-regulated in buds and caulonemal cells. Caulonema are fast growing cells, serve to colonize new substrates and are induced under nutrient deficit. Thus, genes involved in vesicular trafficking and autophagy could acquire an important role to sustain rapid growth and maintain growth under stress conditions. In *A. thaliana, FYVE1* (*FREE1*) is ubiquitously expressed and has important roles in intracellular trafficking and autophagy. AtFYVE1 binds to PtdIns3*P* through its FYVE domain and to ubiquitin, and interacts with Vps23 allowing its incorporation into the ESCRT-I complex required for vacuolar sorting of ubiquitinated membrane proteins (Gao et al. 2014). Also, it also interacts with SH3P2 and associates with the PI3K complex to regulate the autophagic degradation in plants (Zhuang and Jiang 2014).

*PpPRAF2* is up-regulated in caulonema but also in buds (gametophore apical cells) which are meristematic cells. A *M. truncatula* PRAF, *MtZR1*, with 51,4% similarity with PpPRAF2, is mainly expressed in the root tip, the meristems of emerging lateral roots and during nodule development (Hopkins et al. 2014). Additionally *MtZR1* has been suggested to be involved in the development of root and nodules (Hopkins et al. 2014). Several lines of evidence indicate that a root-like developmental program is operational in the nodule meristem, where *WOX5* and *PLT1-4* are essential regulators (Franssen et al. 2015). Interestingly, analysis of *PpPRAF2* promoter reveals binding sites for AP2-type TFs such as *PLT1* and *AIL7* (Supplemental table 7), which are essential regulators of gametophore apical cells in *P. patens* (Aoyama et al. 2012), raising the possibility that *PpPRAF2* could have a role to promote the acquisition of the 3D fate necessary for leafy shoot formation.

*PpALFY3* belongs to proteins from class VII. A mammalian protein with a similar domain structure to class VII is ALFY (Autophagy Linked FYVE protein), a large scaffolding protein required for degradation of protein aggregates by selective macroautophagy (Filimonenko et al. 2010). ALFY interacts directly with the autophagy receptor p62 through its PH and BEACH domains, to Atg5 through the WD40 domain and with PtdIns3*P*-containing membranes through its FYVE domain (Filimonenko et al. 2010, Simonsen et al. 2004).

### *PpSNXs* are PHOX members mainly express during development of *P. patens*

*PpSNX1*, *PpSNX2*, *PpSNX4c* and *PpPX2* were *PHOX* genes mainly up-regulated in caulonemata and buds. Orthologs of PpSNX1 and PpSNX2 in *A. thaliana*, AtSNX1, AtSNX2a and AtSNX2b, are components of the SNX subcomplex of the retromer, but the triple mutant in *A. thaliana* displays only minor developmental defects under standard growth conditions (Pourcher et al. 2010). However, it has been postulated that SNX proteins may represent a potential stress-response trafficking module in plant cells, required upon an enhanced demand of specific protein recycling (Brumbarova and Ivanov 2016). Co-localization analyses showed that AtSNX1 positive-endosomes are involved in the trafficking and recycling of several PM proteins such as PIN2 efflux auxin carrier under high temperatures (Jaillais et al. 2006, Hanzawa et al. 2013), or the iron transporter IRT1 in the regulation of iron homeostasis upon iron deficiency (Ivanov et al. 2014). Homologs of *PpSNX4s* in *A. thaliana* have also been highly expressed in young tissue, namely young seedlings, in young leaves, root tips, and developing embryos, and AtEREX1 is involved in the biosynthetic trafficking to vacuoles as an effector of RAB5 (Sakurai et al. 2016). So far, the role of these proteins in *P. patens* is unknown.

### *PpFYVE* and *PpPHOX* exhibit subtle changes at the transcriptional level upon abiotic stresses suggesting a posttranslational control

Transcriptional changes in response to abiotic stresses for *PpFYVE* and *PpPHOX* were observed, with a general down-regulation pattern under the conditions tested (Figure 8). Small changes in gene expression were also reported for *FYVE* genes from *O. sativa* under abiotic stress even at earlier time points (15 min, 30 min, 1h), in addition for genes encoding AtPPI-modifying enzymes under different stresses (Heilmann 2016, Xiao and Shaw 2016). Although our data demands an expression analysis at earlier time points (data analyzed here 30 min and 4h) to detect rapid and transient changes in transcription, it is plausible that these proteins are mainly controlled at a posttranslational level. In addition, the down-regulation of *FYVE* and *PHOX* genes under abiotic stress conditions in contrast with the pattern observed under development also suggest a mechanism of stress-response activation which generally comes at the expense of plant growth or growth–defense tradeoff (Bechtold and Field 2018). Analysis of *PpFYVE* and *PpPHOX* promoters revealed several elements associated with stress responses to low temperature and drought, defense and hormones such as salicylic acid, methyl jasmonate and abscisic acid which participate in stress responses (Supplemental Table 5). The *P. patens* genome encodes all components of the core ABA signaling pathway identified in angiosperms, and interestingly several *PpFYVE* and *PpPHOX* genes which contains ABRE, DRE and MBS elements in their promoters are upregulated in the *anr* mutant, which fails to respond to ABA in comparison to the WT when treated with ABA or dehydration (Figure 6, (Stevenson et al. 2016)). Particularly in the WT, *PpFAB1*, *PpFYVE5*, *PpPRAF2* and *PpALFY1-4* were the genes with a down-regulation pattern under cold, drought and salt. Down-regulation of *MtMZR1* (PRAF) was also observed by drought and salt treatments (Hopkins et al. 2014). In similar trend, all *PpPHOX* were down-regulated under the conditions evaluated and up-regulated in the *anr* mutant, with exception of *PpSNX4c* which was up-regulated by ABA, salt, cold and dehydration treatments. Therefore, *PpSNX4c* is a robust candidate in regulating abiotic stress in *P. patens*.

### Final conclusions

We have identified 13 and 9 genes coding for FYVE and PHOX proteins, respectively in *P. patens*. PpFYVE proteins resulted in 6 subfamilies previously described (Class I-VI) and we reported an additional group (Class VII) whose FYVE domain was lost during evolution to higher plants, whereas PpPHOX proteins are classified into 5 subfamilies (Class I-IV) previously described in *A. thaliana* with the exception of class V (PLDs). Several *FYVE* and *PHOX* genes have been characterized in *A. thaliana*, but there are still classes whose function is unknown as FYVE proteins from Classes III, V, VI, VII, and PHOX proteins from Classes III and IV. Based on our study, candidate genes were identified in *P. patens* for further functional characterization such as *PpPRAF2*, *PpALFY3*, *PpFYVE4* during the development of tip growing cells and buds, and *PpSNX4c* under abiotic stress where exciting roles may arise. Proteins as FABs, FYVE1 (FREE1), EREX and SNXs are involved in the endocytic pathway; but recently the role of these proteins such as FYVE2 and FYVE1 at the intersection between the conventional endocytic-based membrane trafficking system to the vacuole and autophagy in regulating vacuolar degradation have started to emerge and suggest further studies. Finally, our work reinforces the notion that expanding in the use of other organisms such as, Bryophytes will provide progress in our understanding on fundamental aspects of plant development and evolutionary function of phosphoinositide signaling in plants.

## ADDITIONAL INFORMATION

Supplementary information that accompanies this paper:

**Figure S1.**
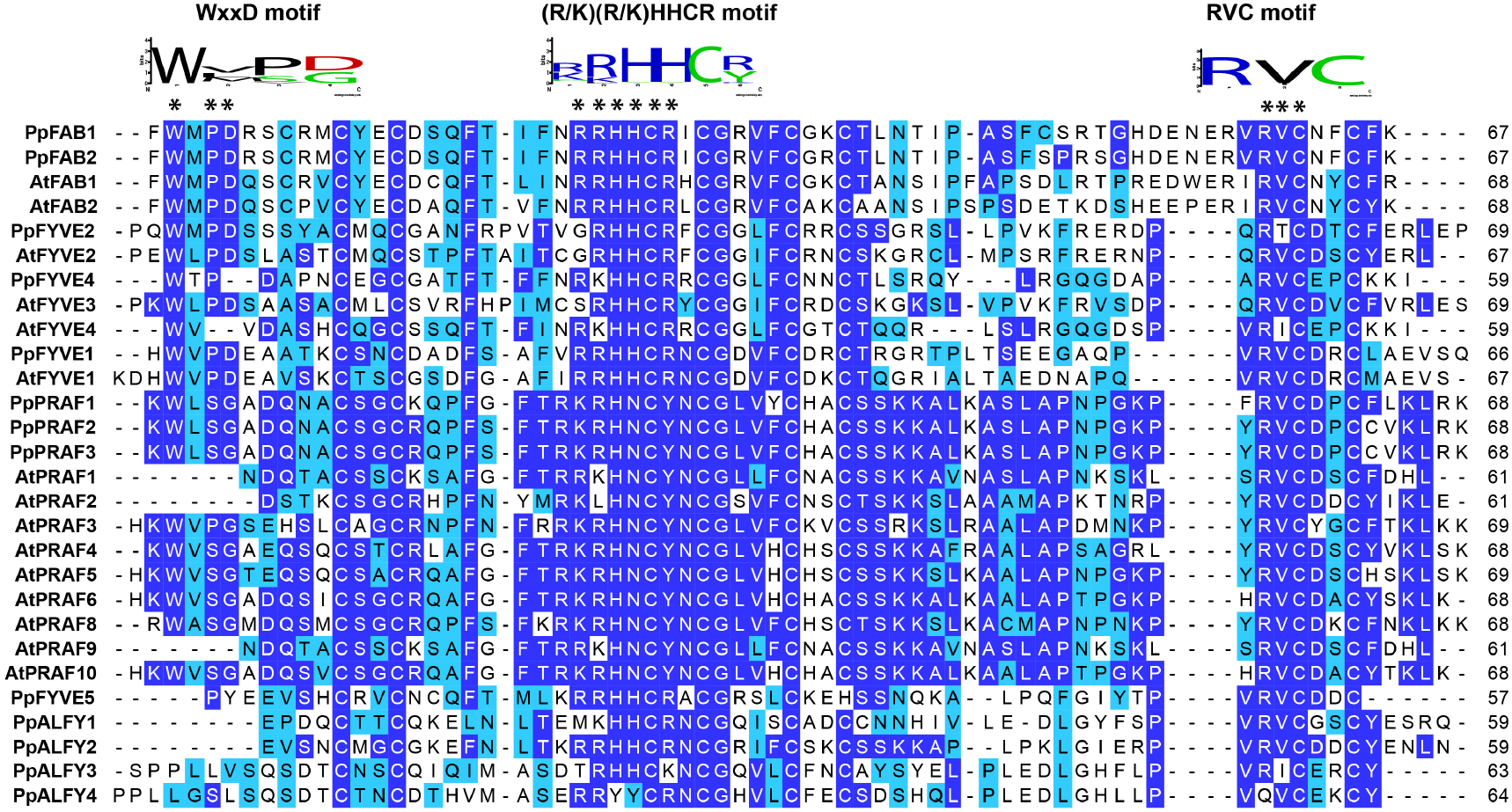
Sequence alignment and motif analysis of the FYVE domain of the repertory of FYVE proteins in *P. patens* and *A. thaliana*.

**Figure S2.**
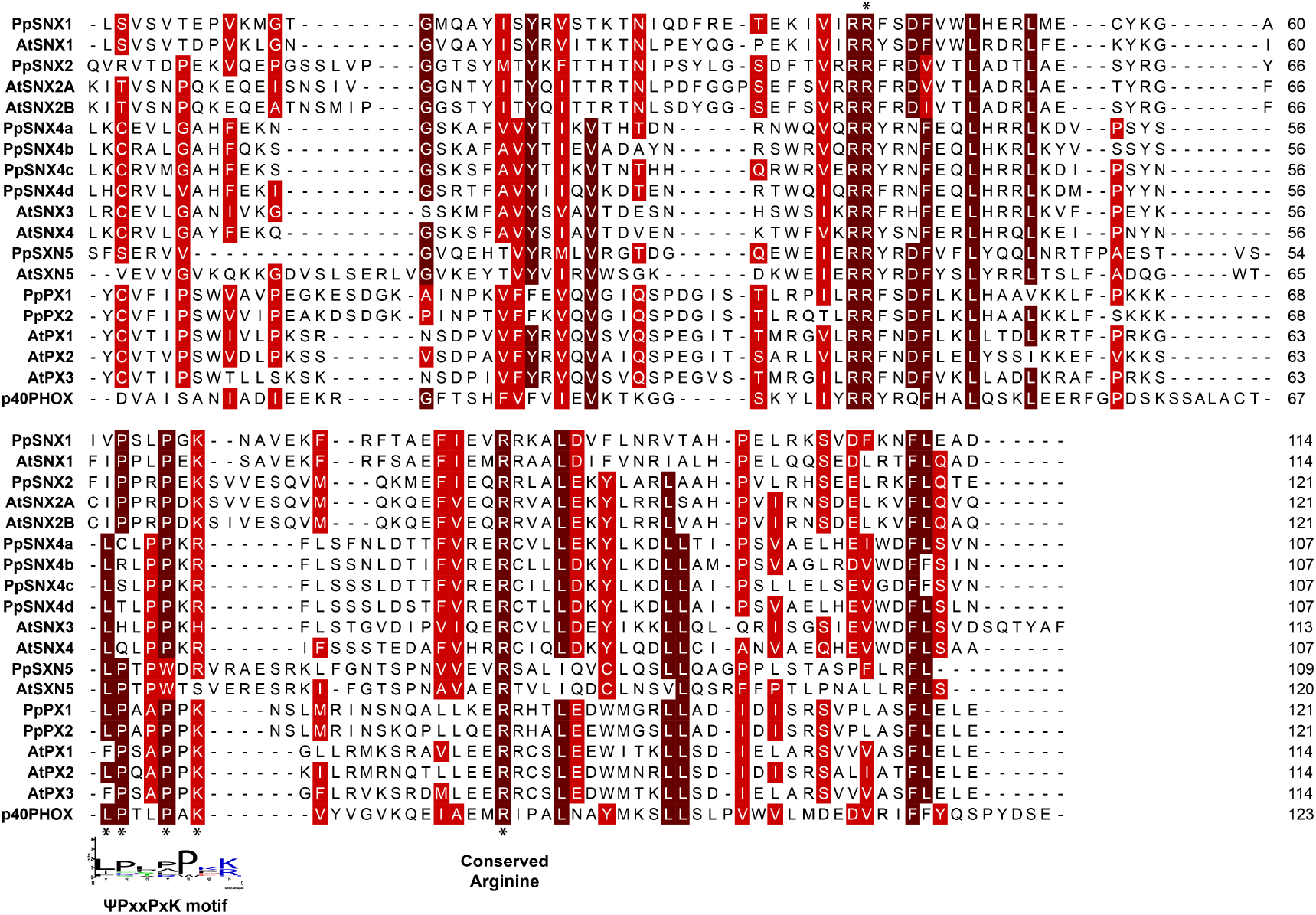
Sequence alignment and motif analysis of the PHOX domain of the repertory of PHOX proteins in *P. patens and A. thaliana*.

**Figure S3.**
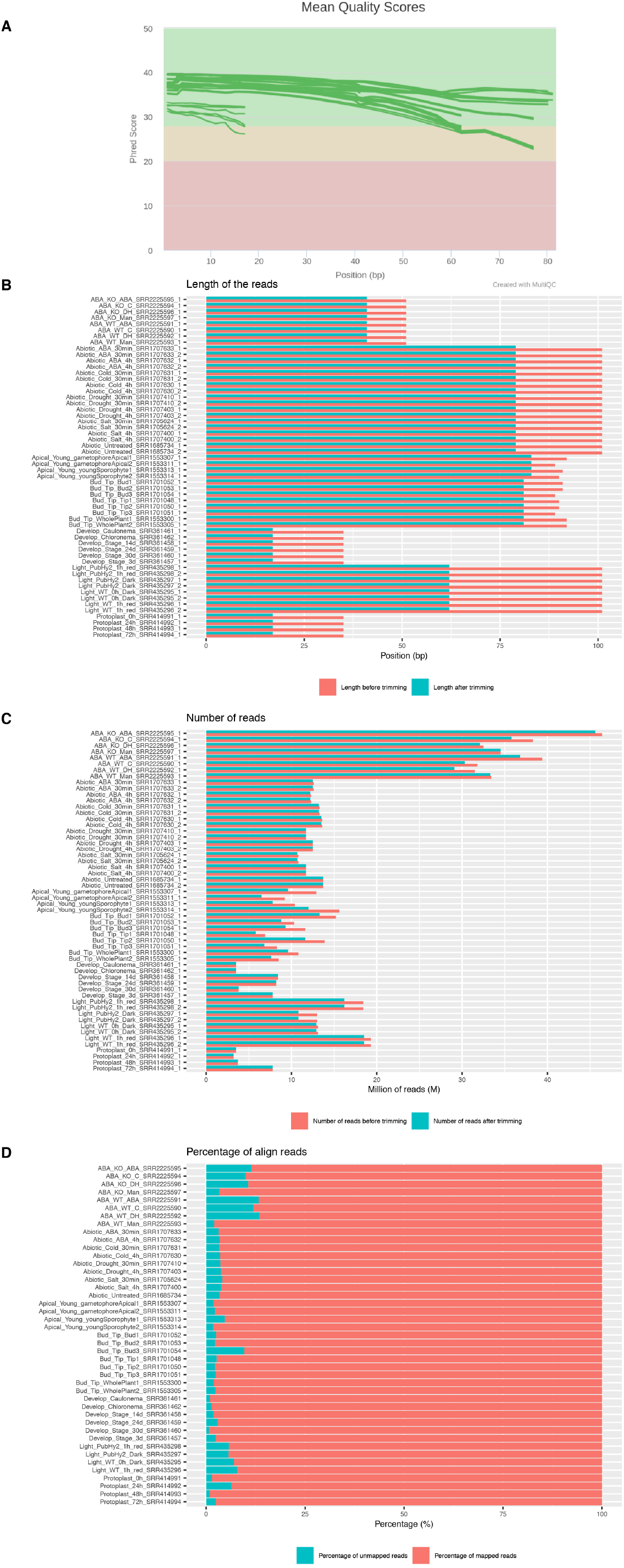
Summary of statistics of the several RNA sequencing studies assessed here. (A) Mean Quality Scores graphic. Lines in green represent each library with quality score > 20 Phred Score. (B) Barplot shows the library fragment size before (pink) and after (blue) trimming. (C) Barplot depicts the number of reads by library before (pink) and after (blue) trimming. (D) Stacked barplot summarizes the percentage of mapped (pink) and unmapped (blue) reads by library from the pseudo-bam files obtained by Kallisto (Bray, Pimentel, Melsted, & Pachter, 2016).

**Table S1-** *FYVE* and *PHOX* genes used in this study.

**Table S2-** *PpFYVE* intron-exon structure.

**Table S3-** *PpPHOX* intron-exon structure.

**Table S4-** Pfam scan PLDs.

**Table S5-** *PpFYVE* and *PpPHOX* promoter cis-acting regulatory elements analyzed with PlantCARE.

**Table S6-** *PpFYVE* and *PpPHOX* promoter transcription factor binding sites analyzed with PlantPAN2.0.

**Table S7-** Scoring tables of the known motif analysis (Homer v4.977) of *PpFYVE* genes.

**Table S8-** Scoring tables of the known motif analysis (Homer v4.977) of *PpPHOX* genes.

**Table S9-** Interaction network analysis of the known motifs of the *PpPHOX* and *PpFYVE* genes in Cytoscape (v3.6.1). (a) Motif-gene interaction network file of *PpPHOX* genes. (b) Node-scoring table of *PpPHOX* net build by Molecular Complex Detection (MCODE) App. (c) Motif-gene interaction network file of *PpFYVE* genes. (d) Node-scoring table of *PpFYVE* net build by MCODE.

**Table S10-** Bioproject list of the transcriptomic studies in *Physcomitrella patens* surveyed on the SRA and GEO databases from the NCBI website.

## ACKNOWLEDGEMENTS

This work was supported by grants from the Agencia Nacional de Promoción Científica y Tecnológica, Argentina (FONCYT-PICT-2016-0497), Consejo Nacional de Investigaciones Científicas y Técnicas (CONICET) and Secretaria de Ciencia y Tecnología (SECYT) Universidad Nacional de Córdoba (UNC). Additional funding was provided by the Portuguese Foundation for Science and Technology (FCT Investigator IF/00169/2015), and UID/MULTI/04046/2019 Research Unit grant from FCT to BioISI.

## AUTHOR CONTRIBUTIONS

L.S., P.A-R., and M.F. designed the study. P.A.-R., M.F, L.S. and T.S. analyzed the data. L.S., A.F. and P.A.-R. wrote the manuscript with valuable input from R.L. All the authors revised and approved the manuscript.

