## Supplementary material for "Evolutionary insights into PtdIns3*P* signaling through FYVE and PHOX effector proteins from the moss *Physcomitrella patens*": Table S2 - FYVE introns _exons.docx

Pp3c17_19810
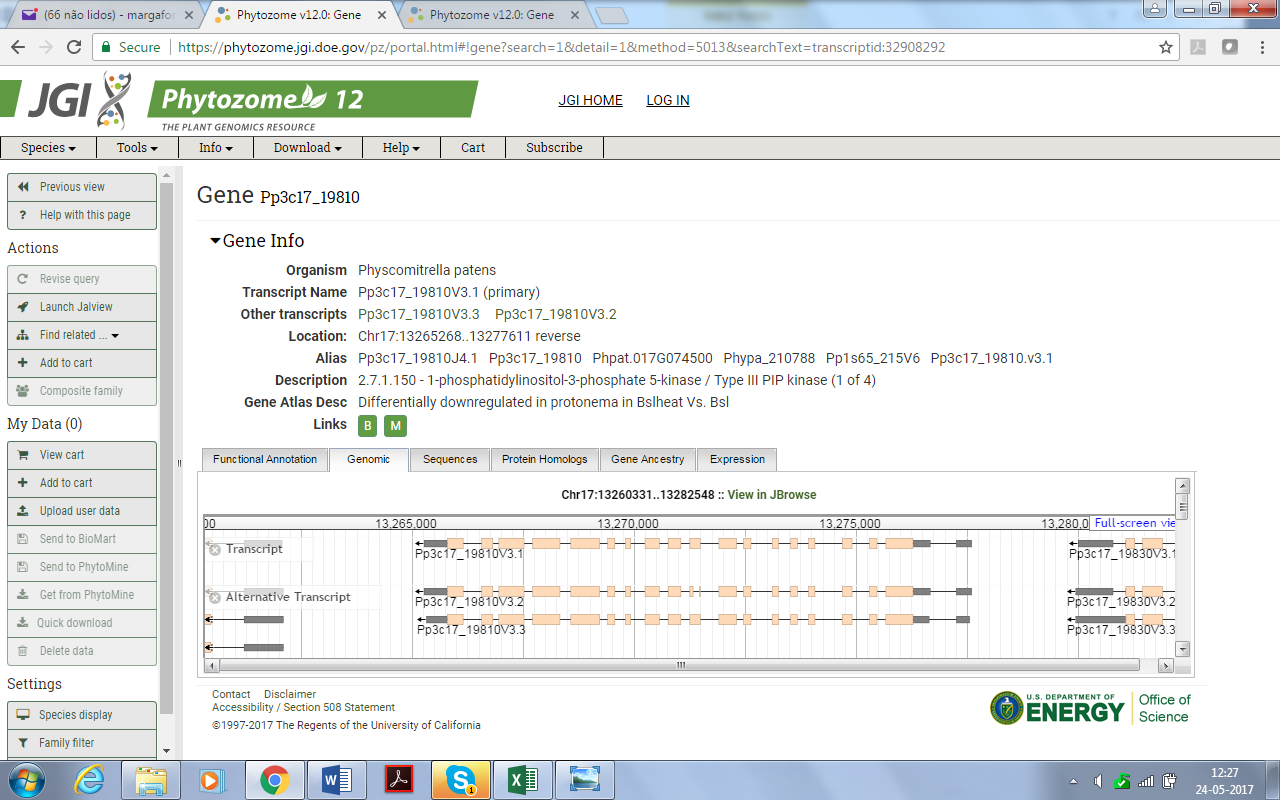

Pp3c14_12410
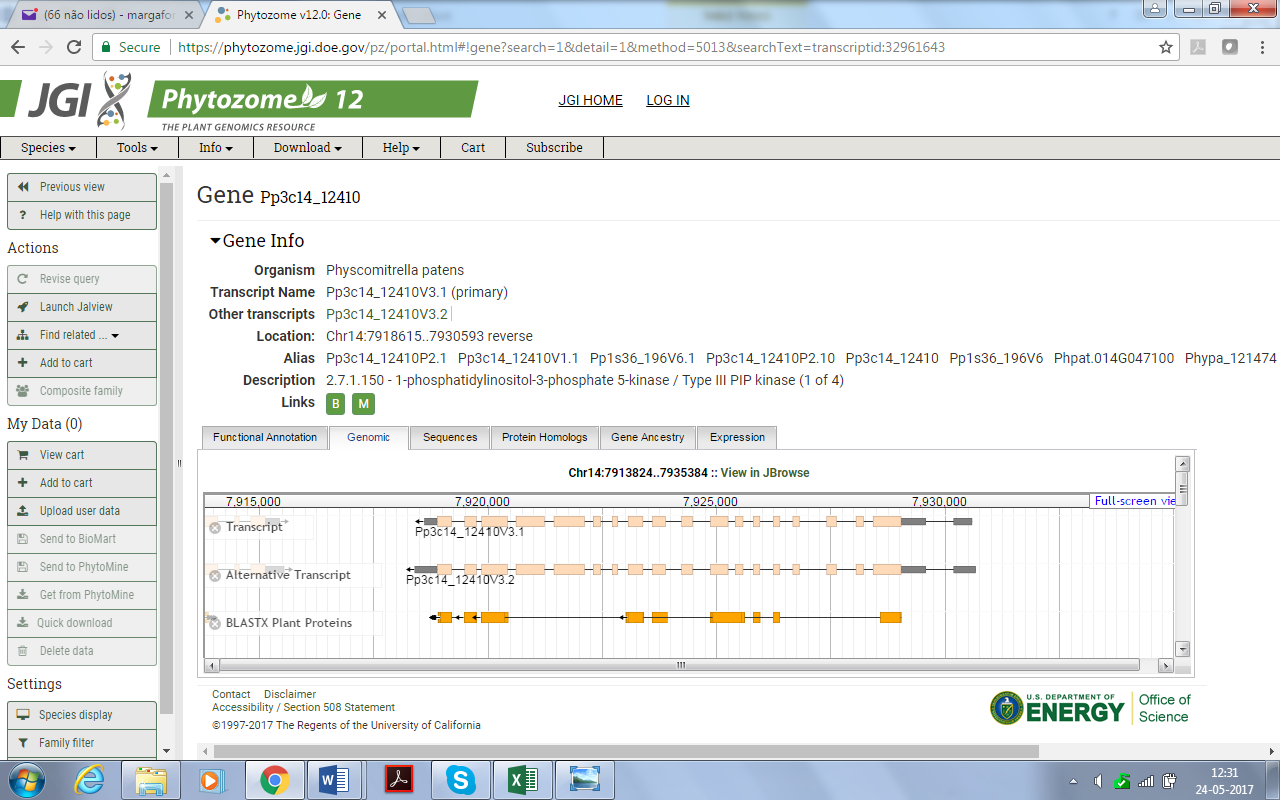

Pp3c21_290
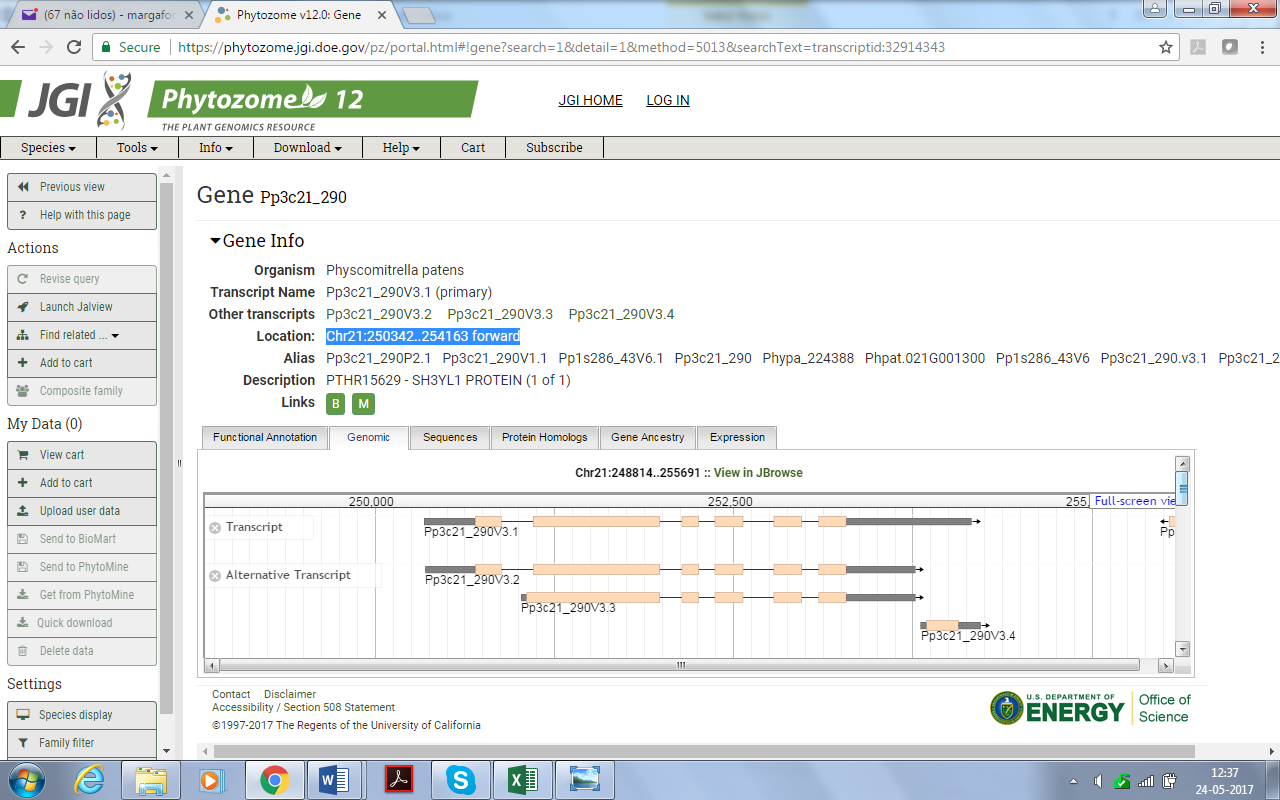

Pp3c16_8350 (for simplicity reasons some versions were not represented in this image)

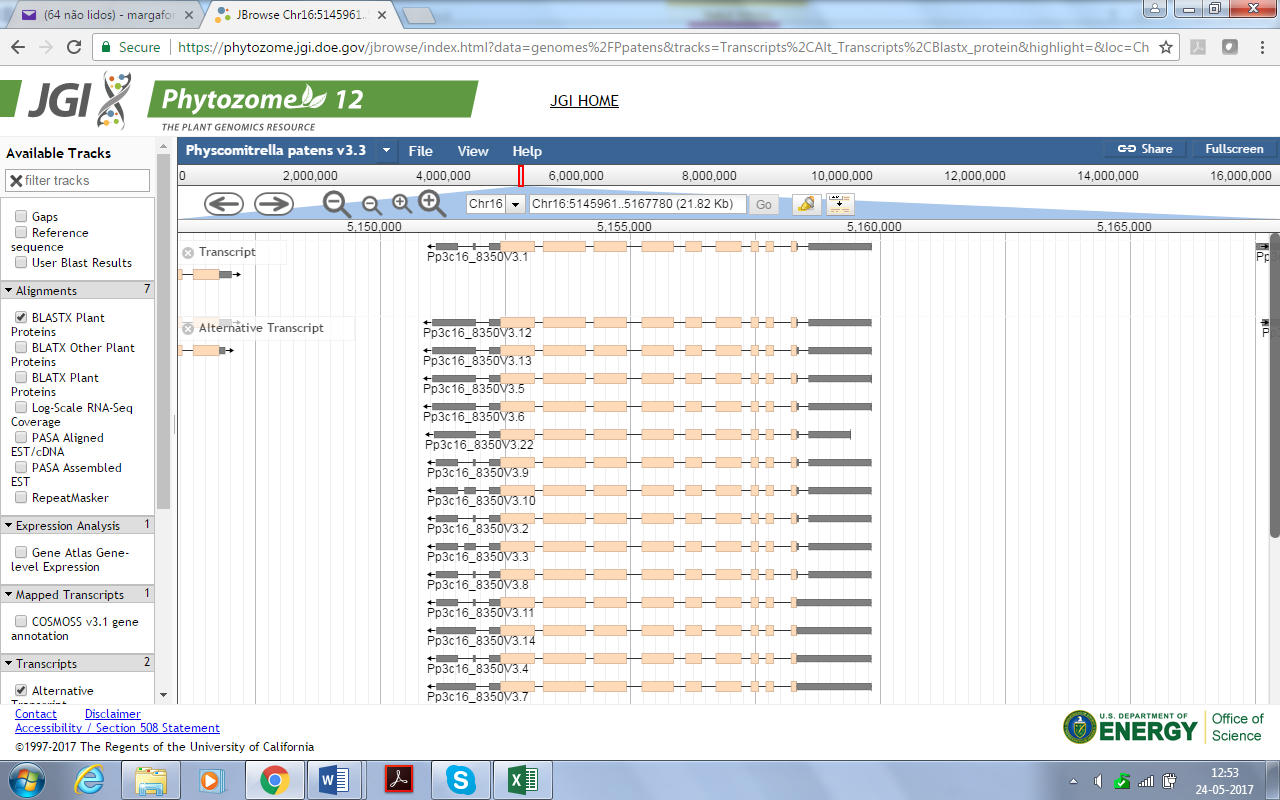

Pp3c14_11260
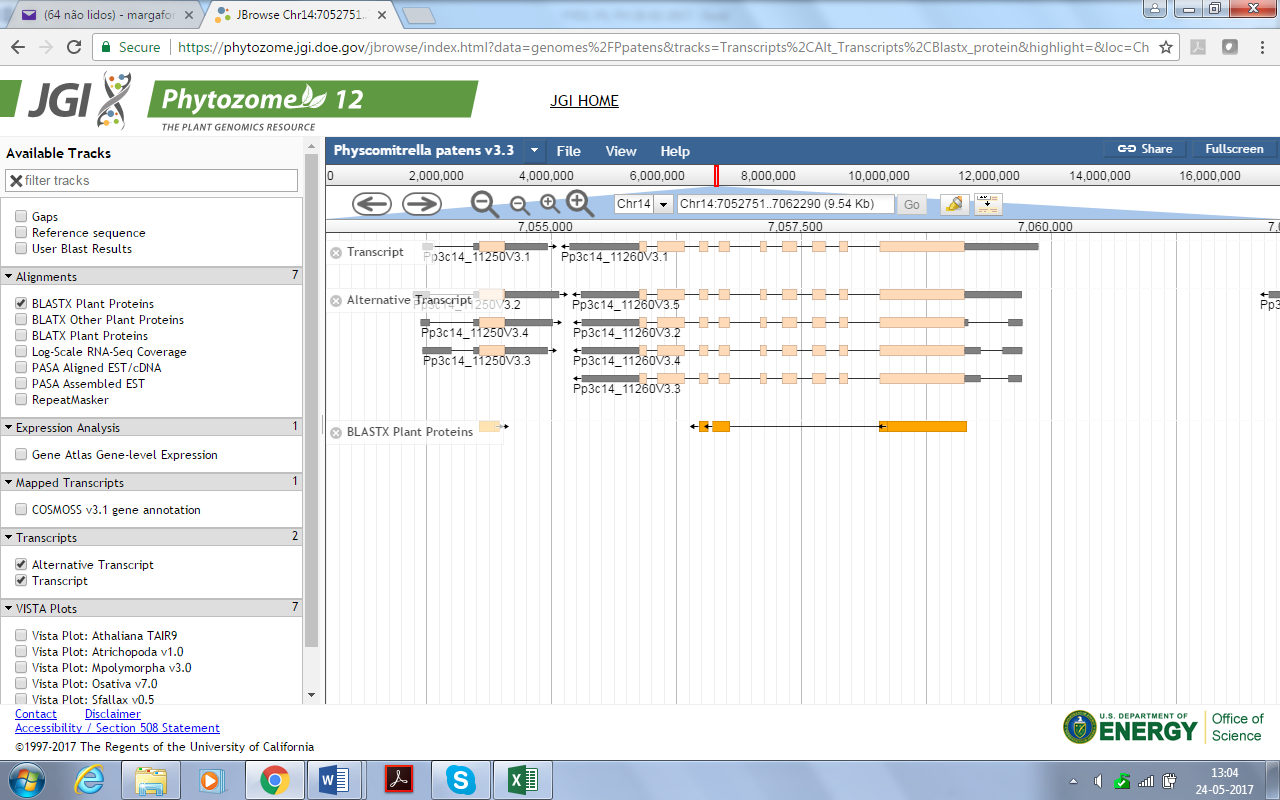

Pp3c10_3893

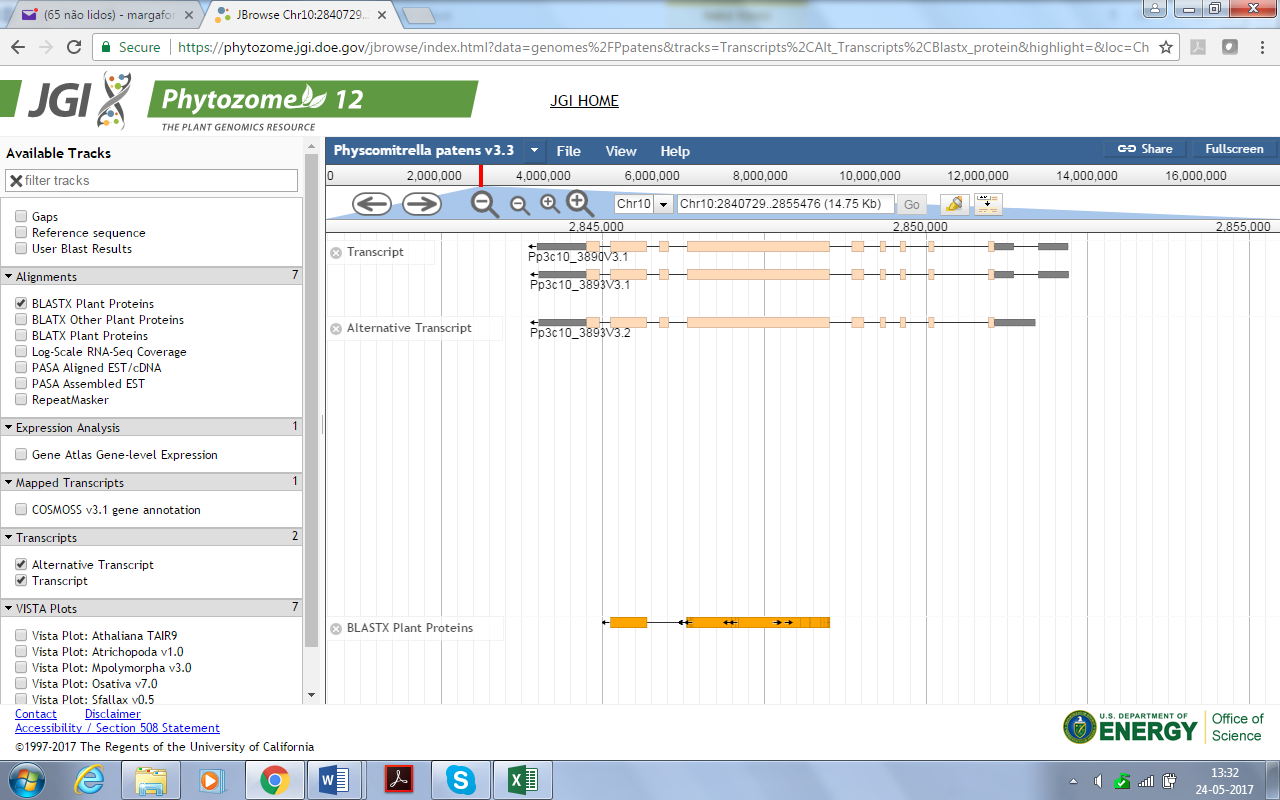

Pp3c10_3890

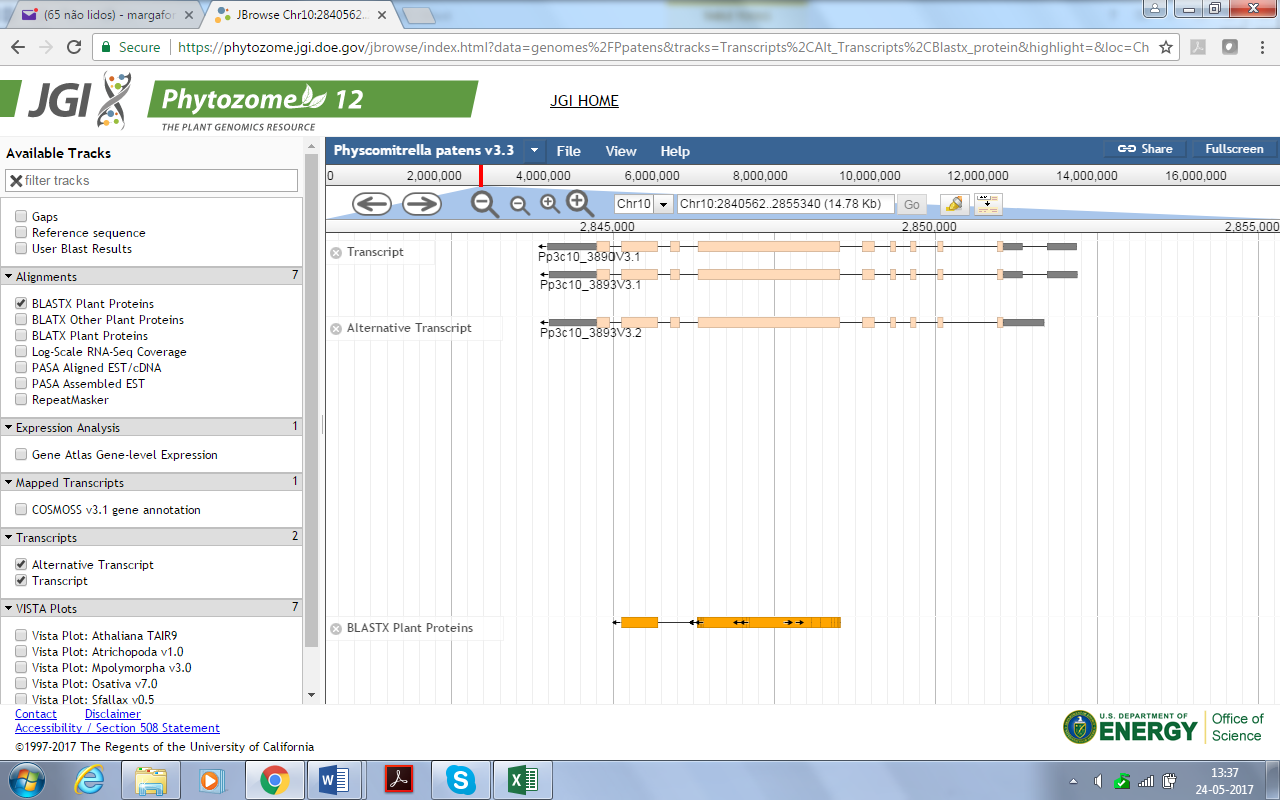

Pp3c14_4760

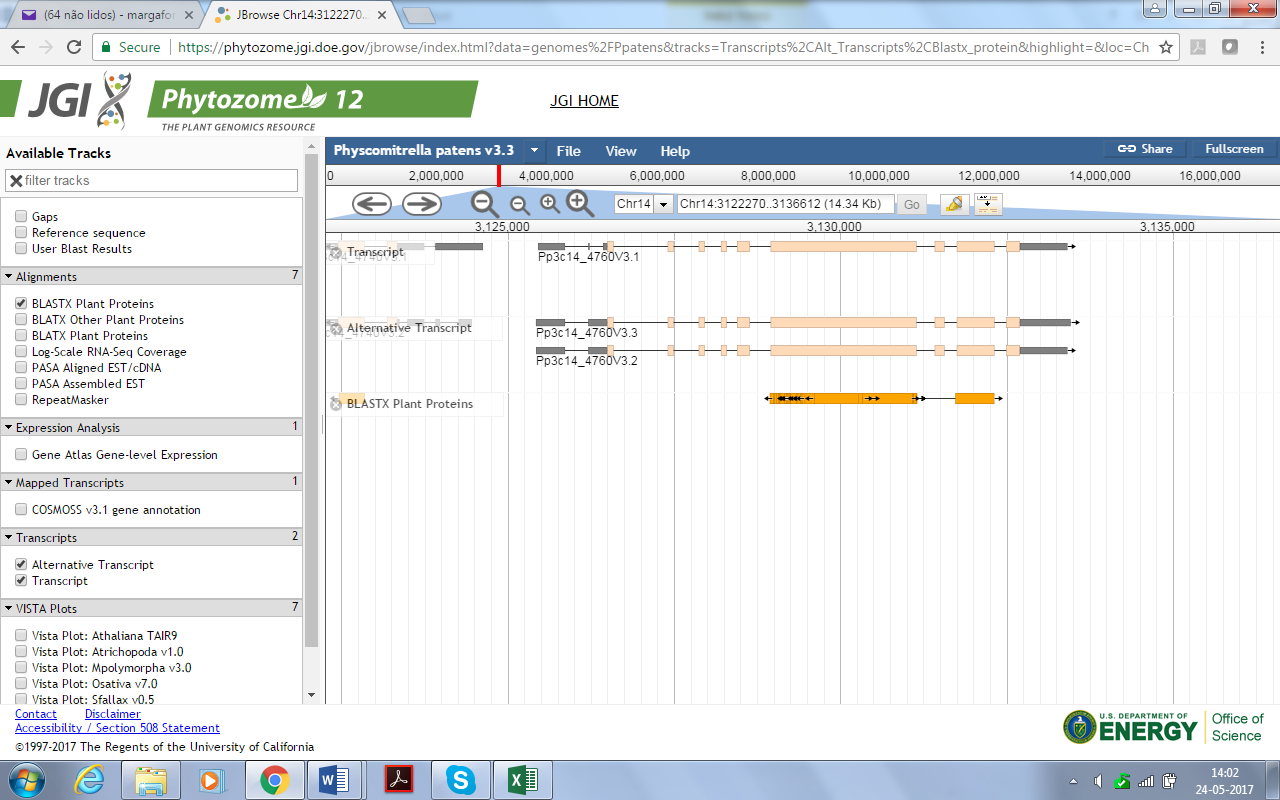

Pp3c14_4640

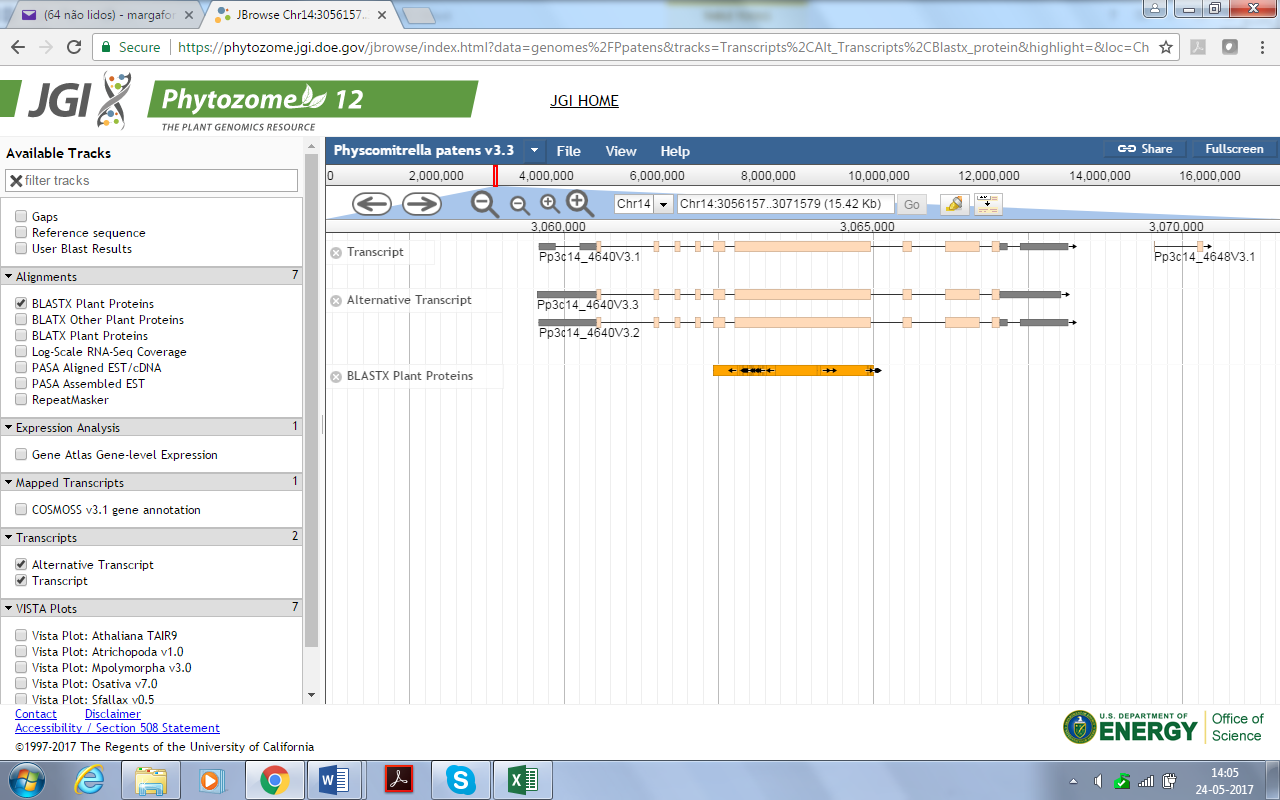

Pp3c18_11010

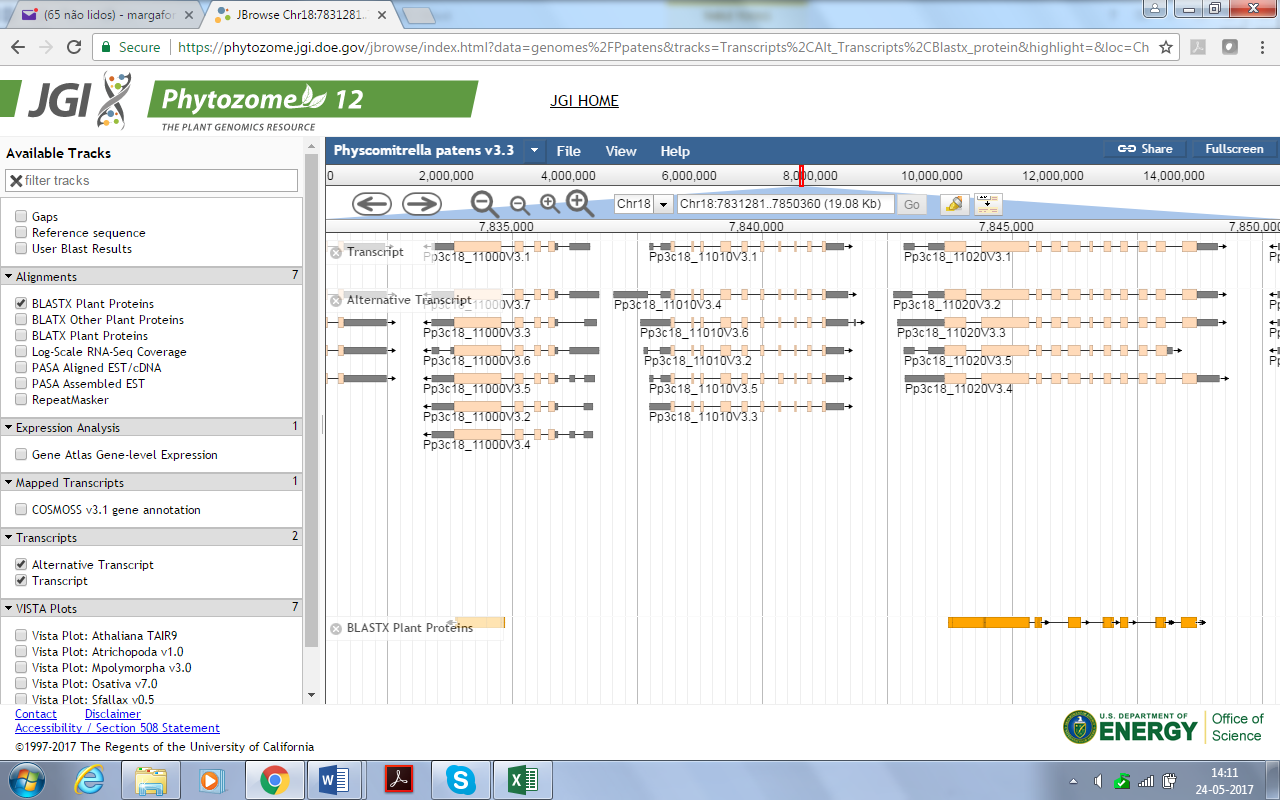

Pp3c6_26100

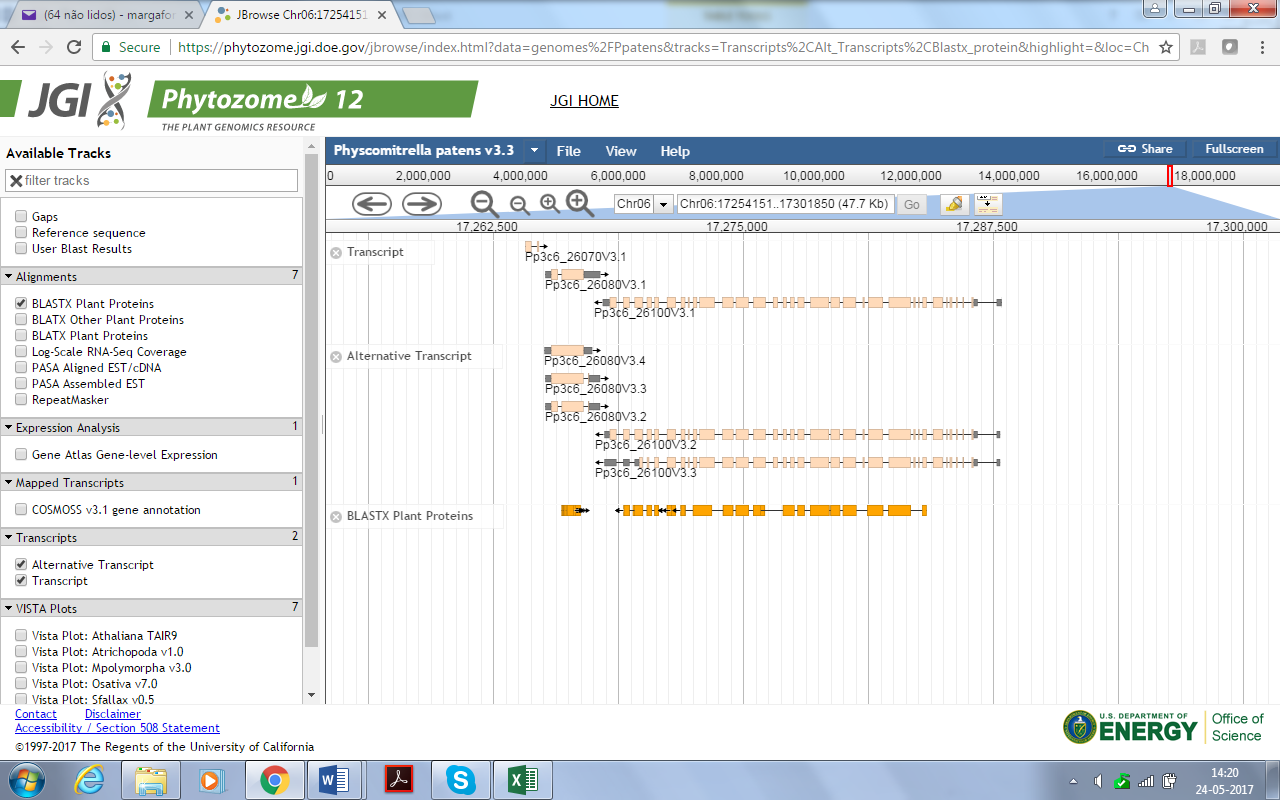

**Pp3c5_2897 and Pp3c5_2890**

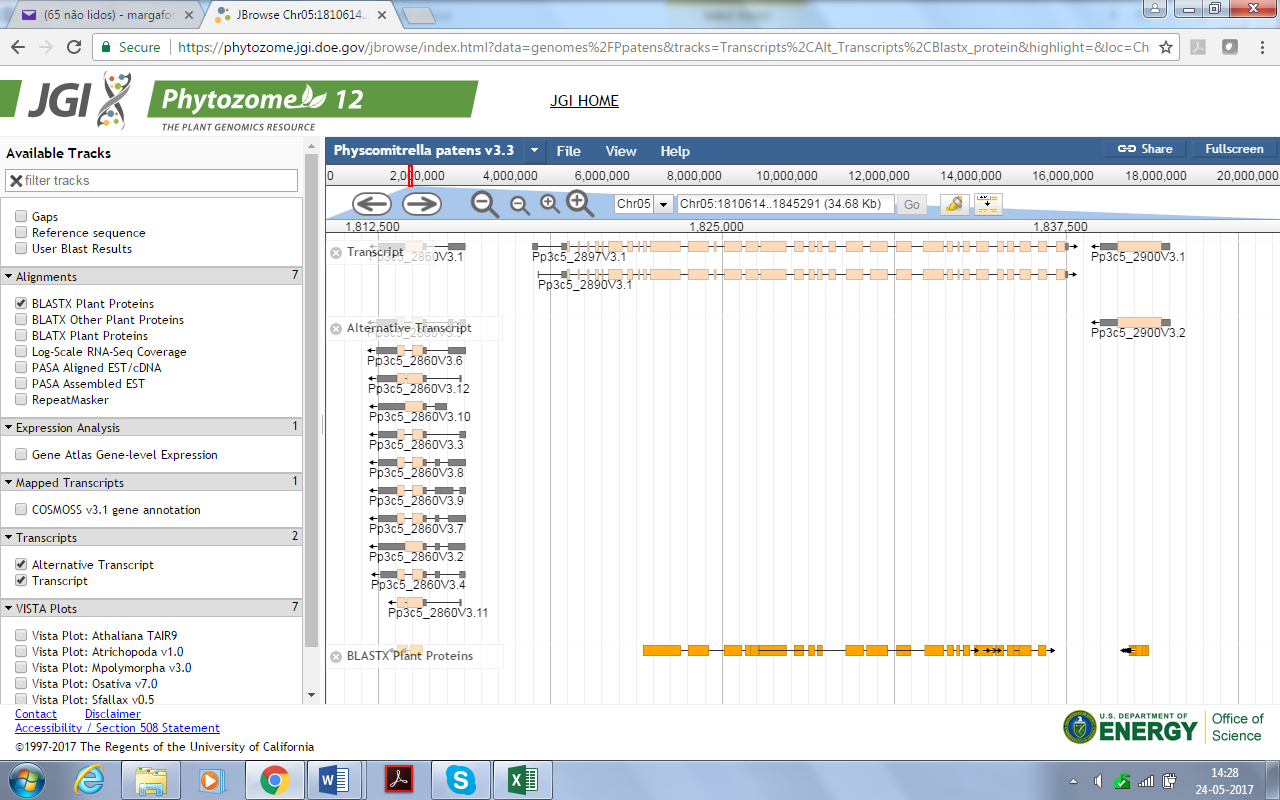

Pp3c16_11700 and Pp3c16_11705

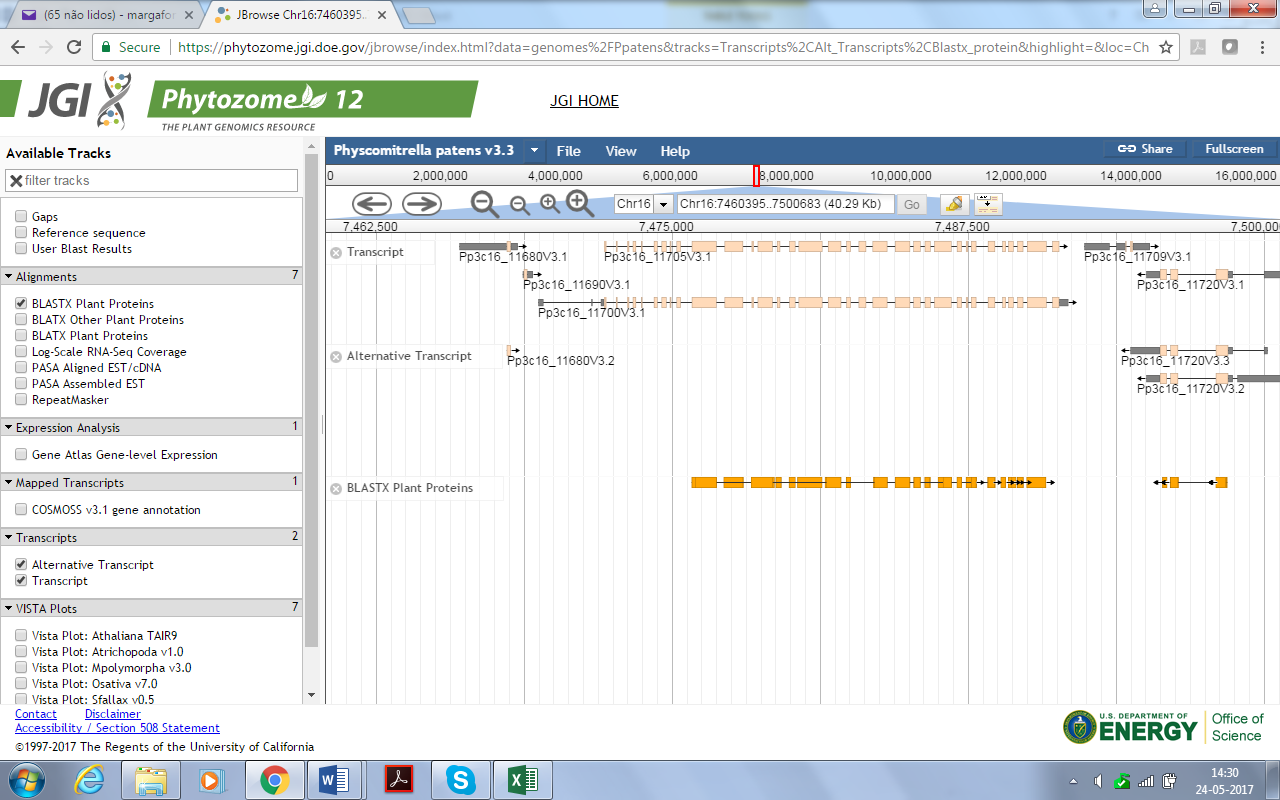

Pp3c6_5706 and Pp3c6_5700

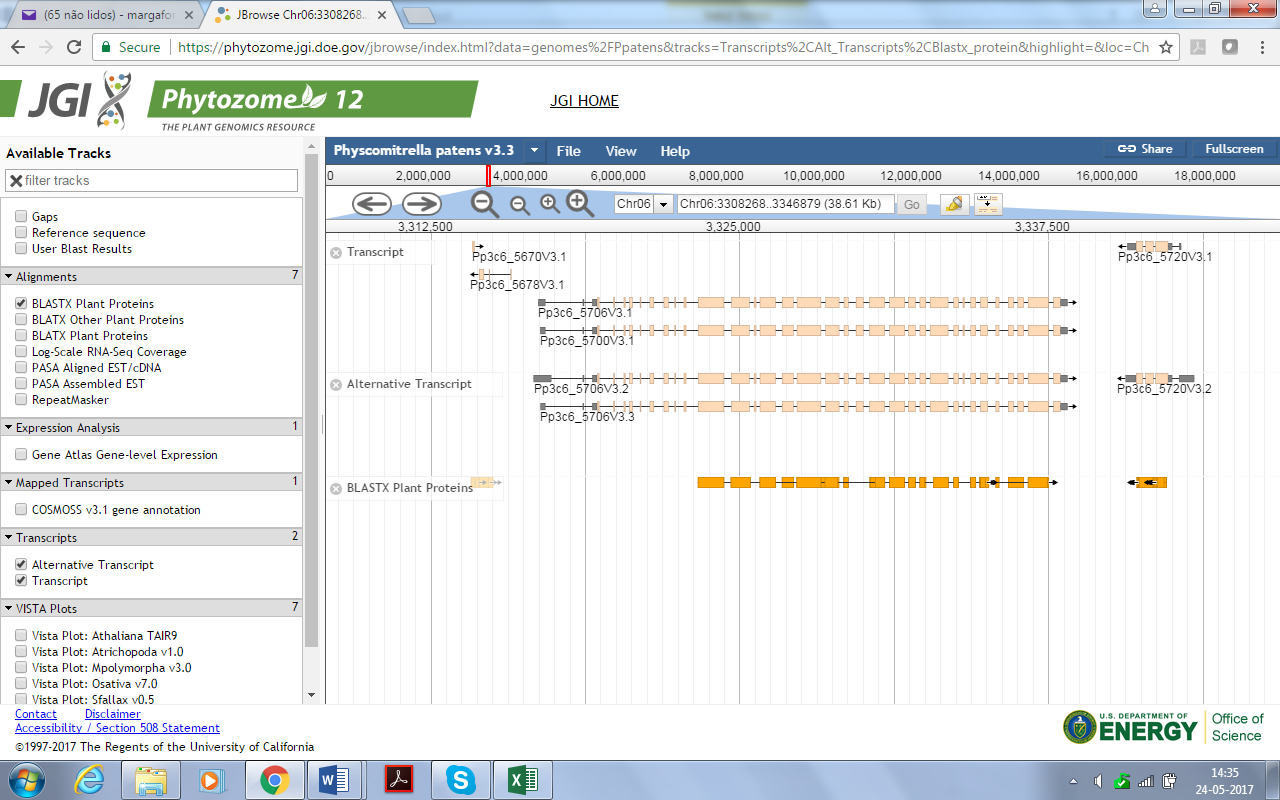
