## Supplementary figures and images for "Evolutionary insights into PtdIns3*P* signaling through FYVE and PHOX effector proteins from the moss *Physcomitrella patens*"

### Figure S1- Fyve_domain.pdf

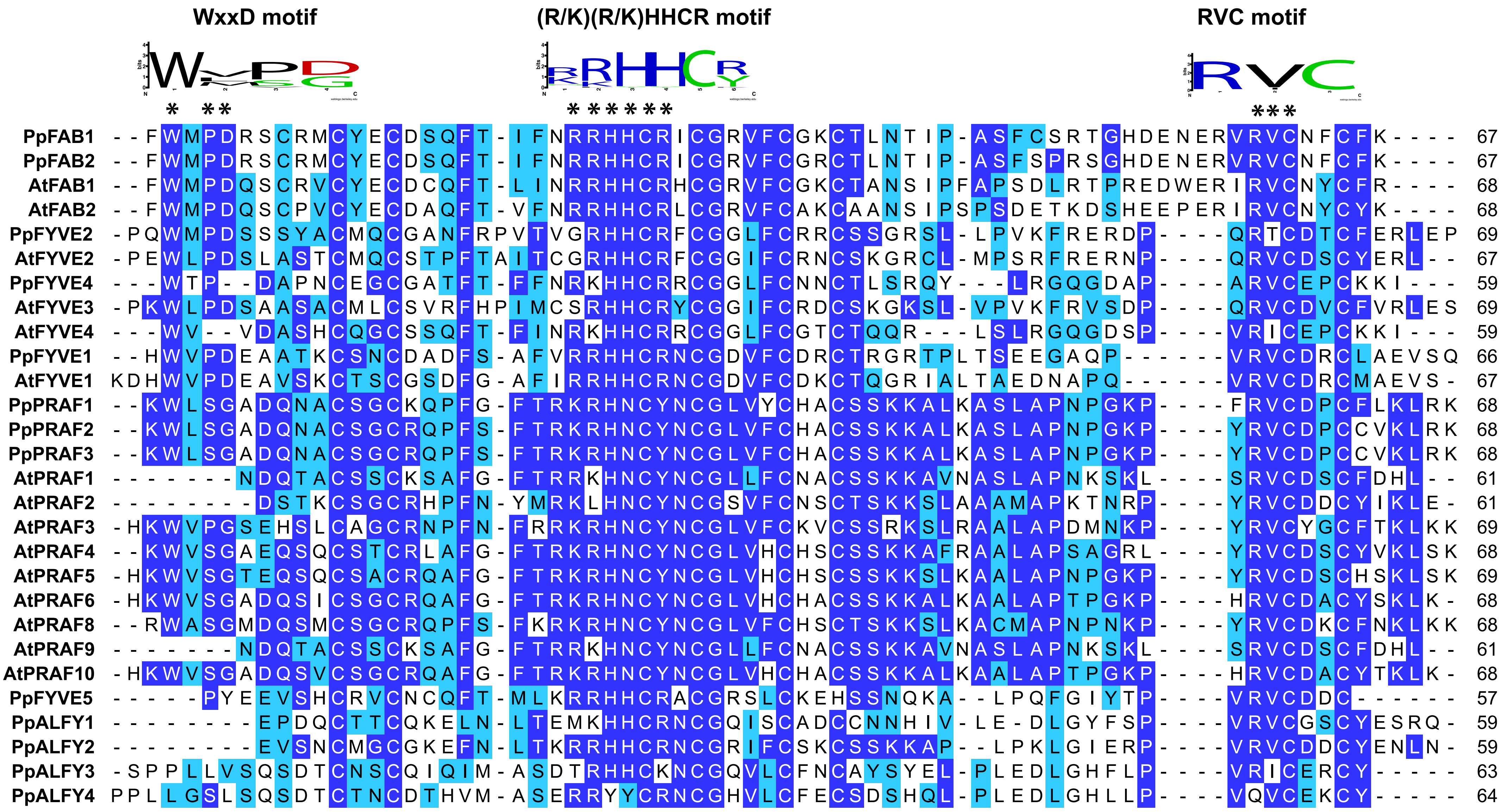

### Figure S2- PHOX domain.pdf

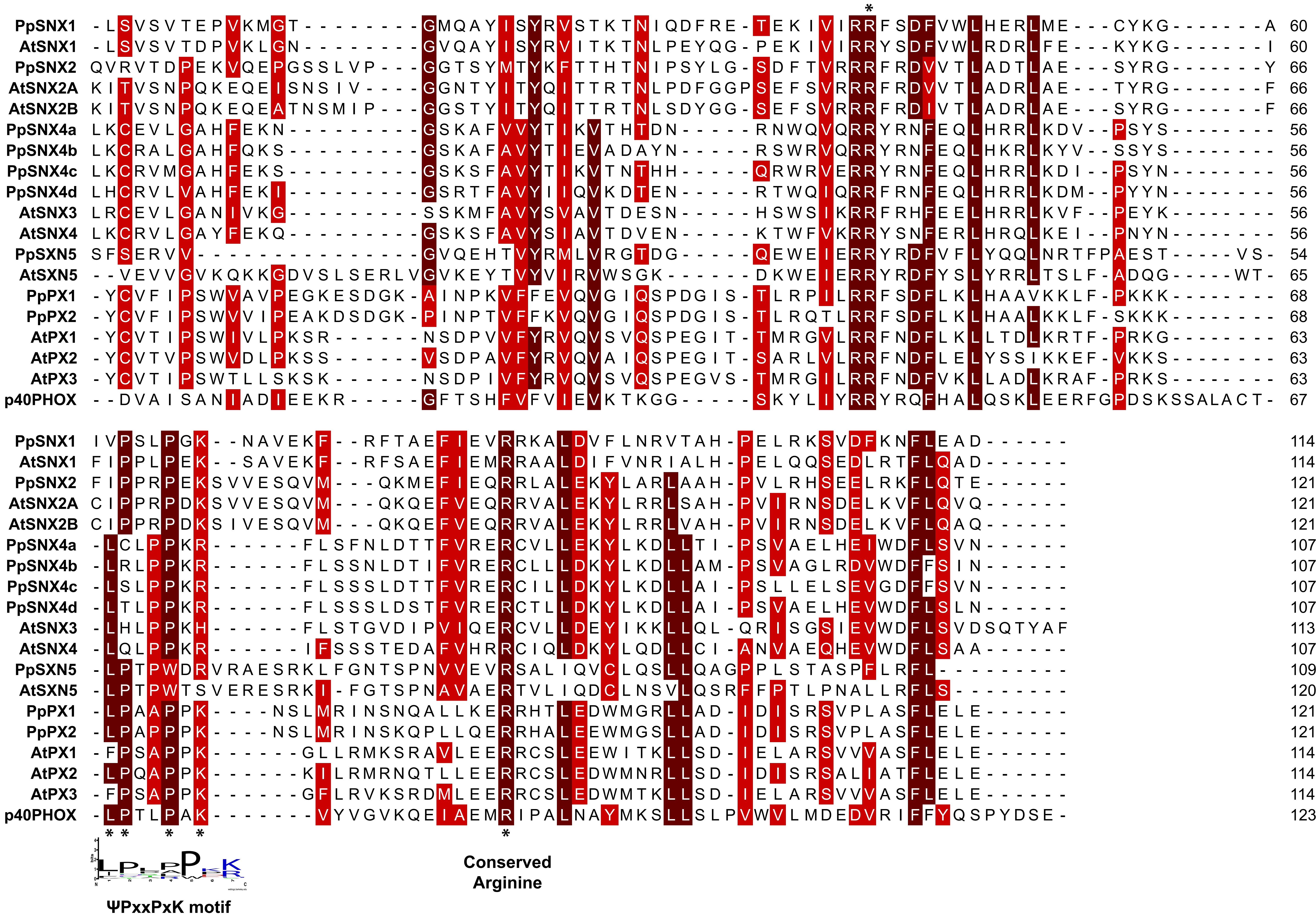

### Figure S3.pdf

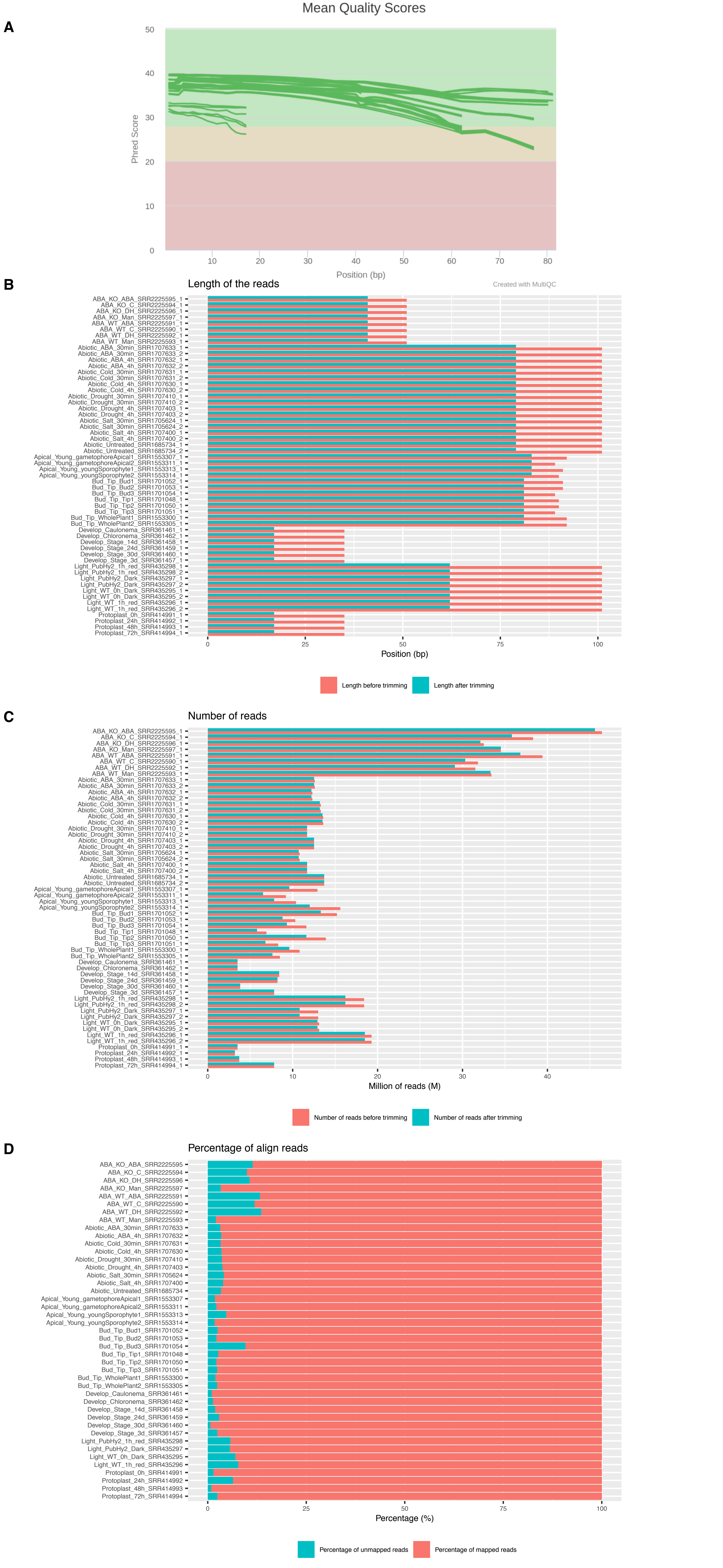

### Table S3- PHOX introns _exons.docx

**CLASS I**

Pp3c1_14920

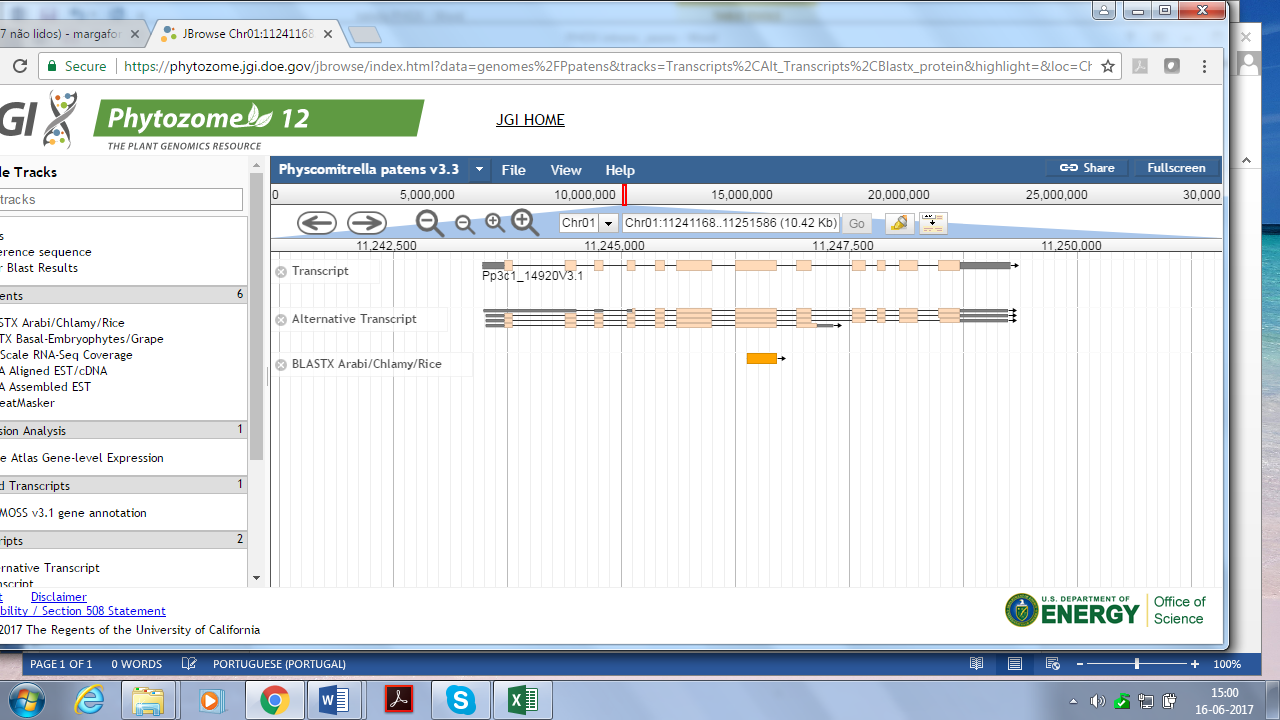

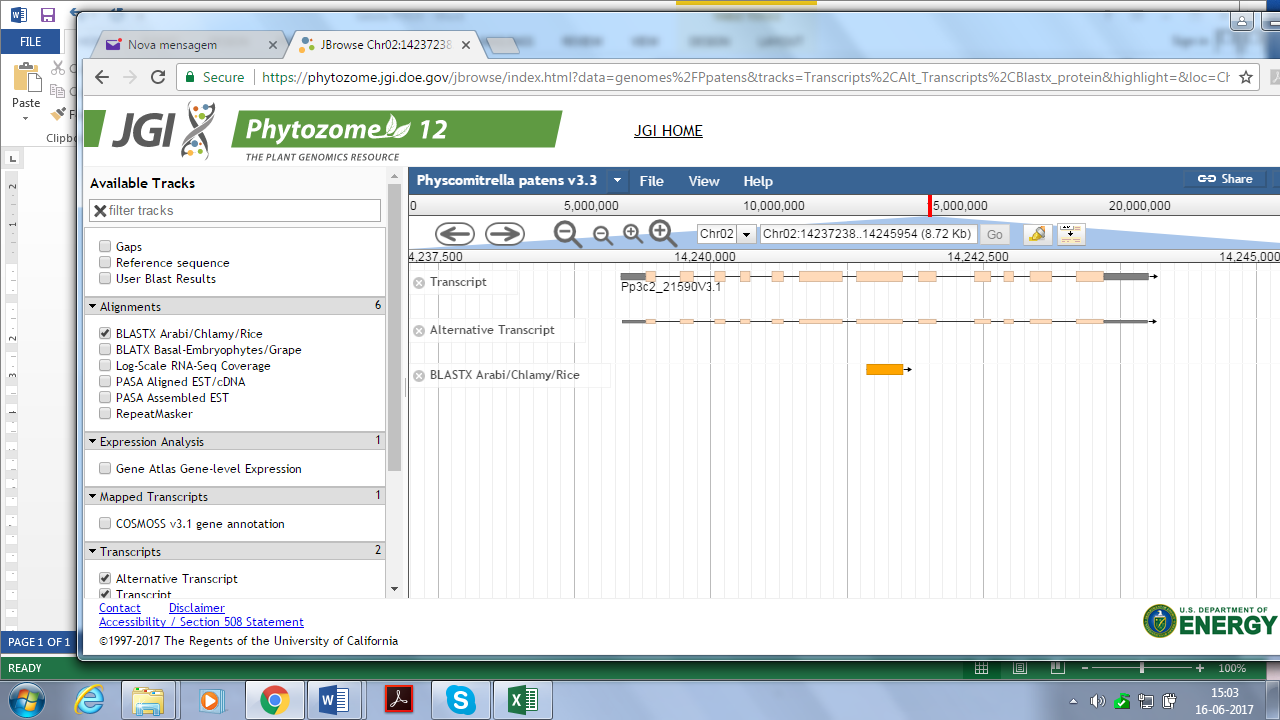
Pp3c2_21590

Pp3c2_34810

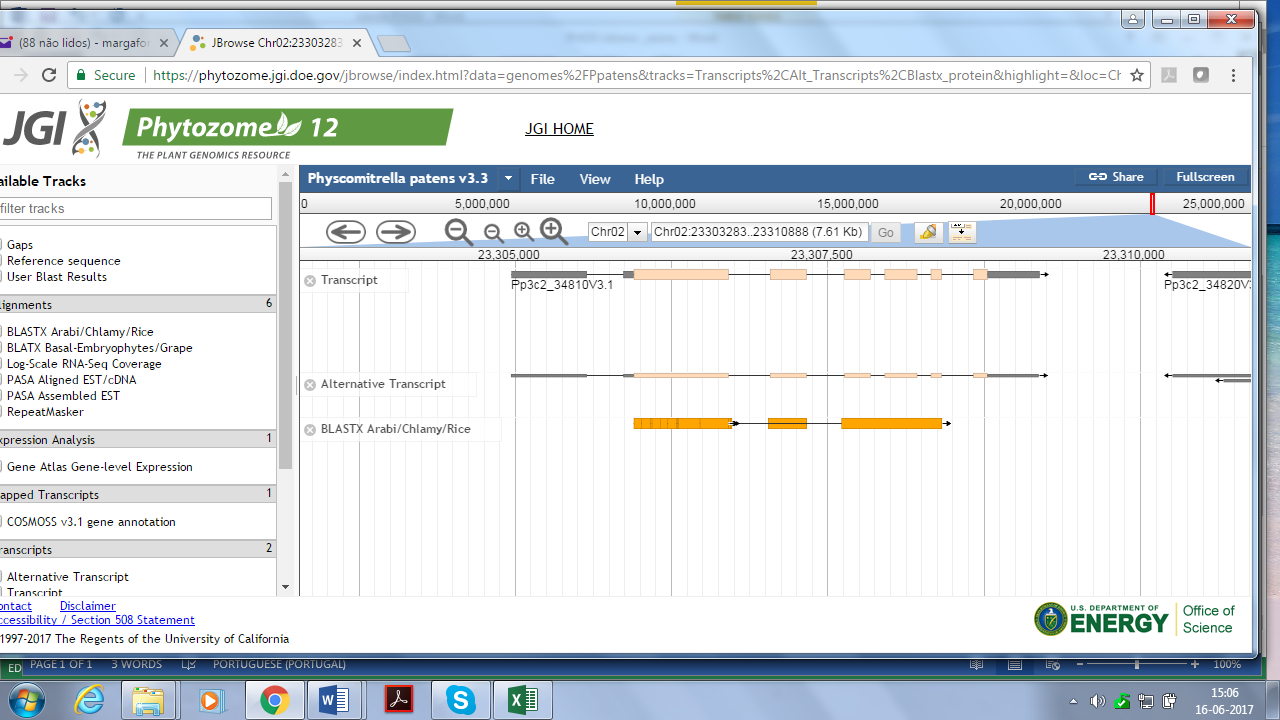

Pp3c4_27410

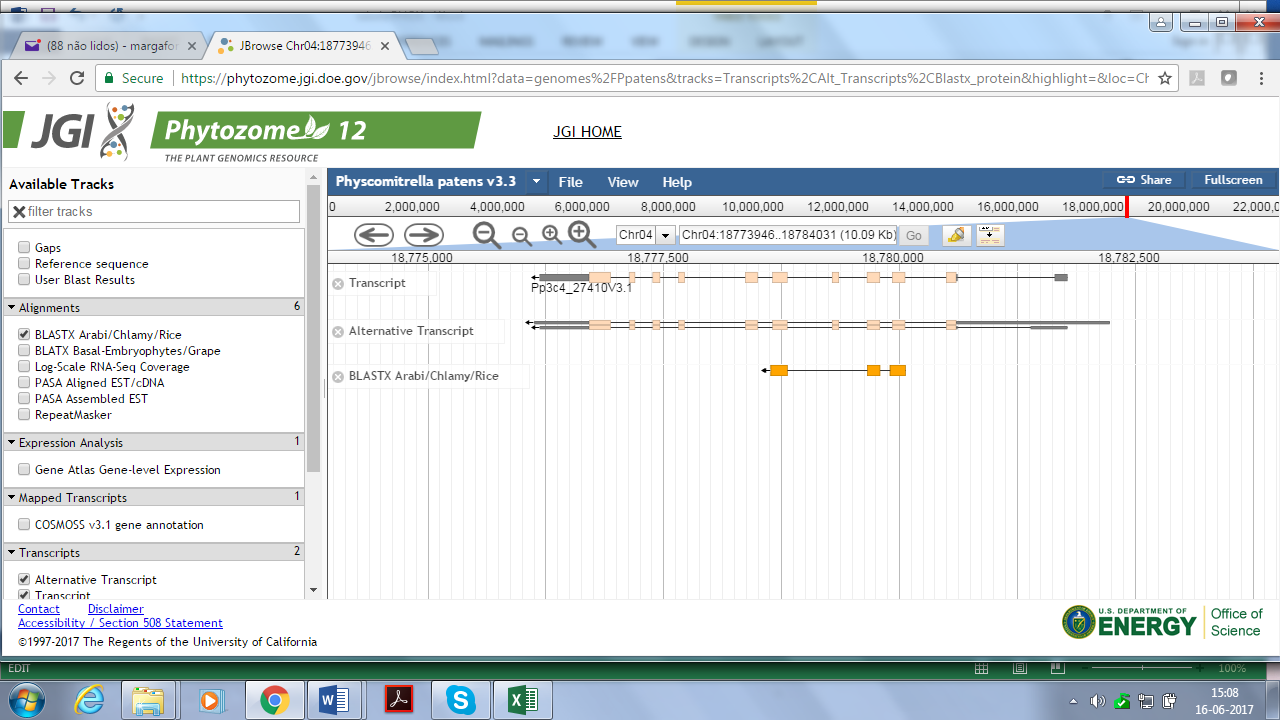

**CLASS II**

Pp3c26_1280

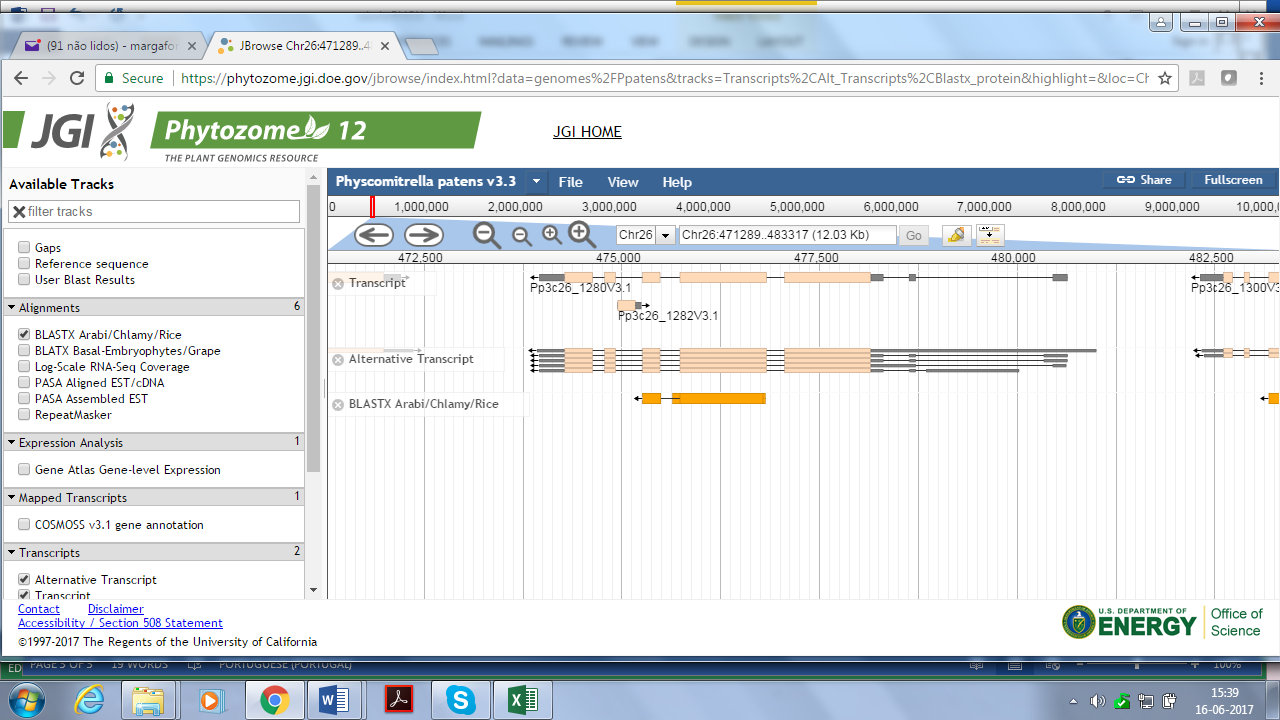

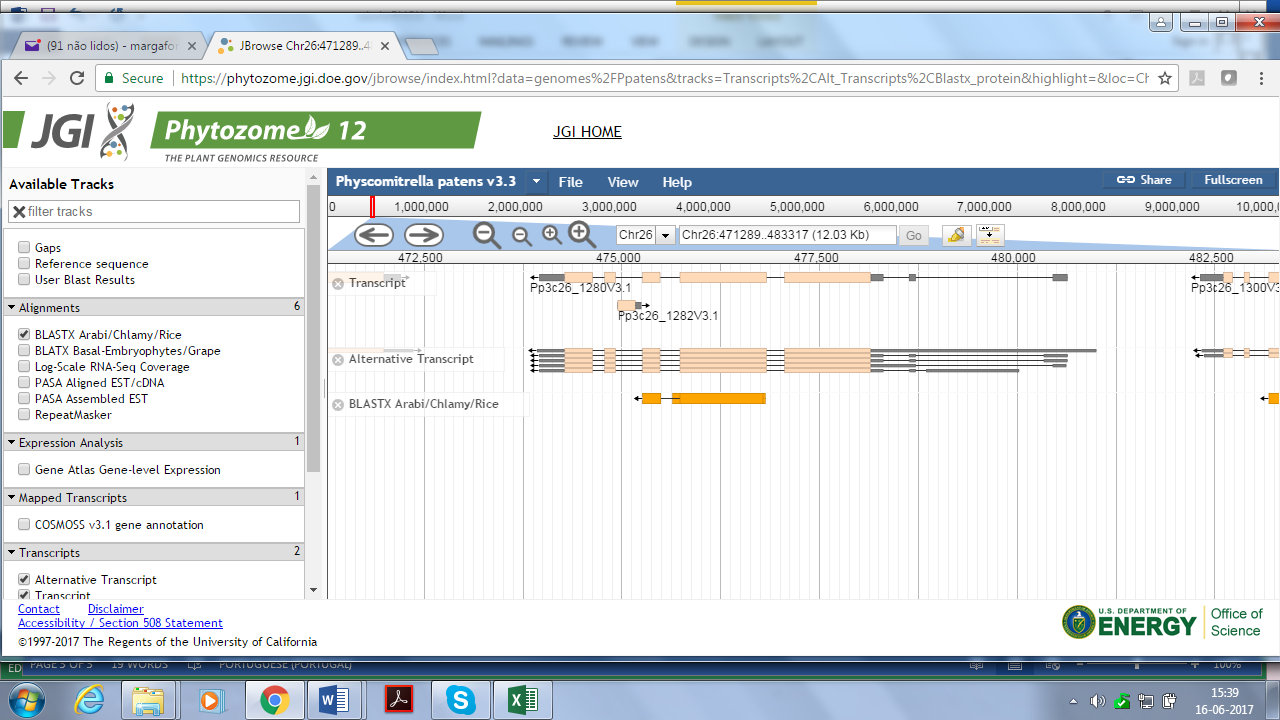

**CLASS III**

Pp3c17_21440

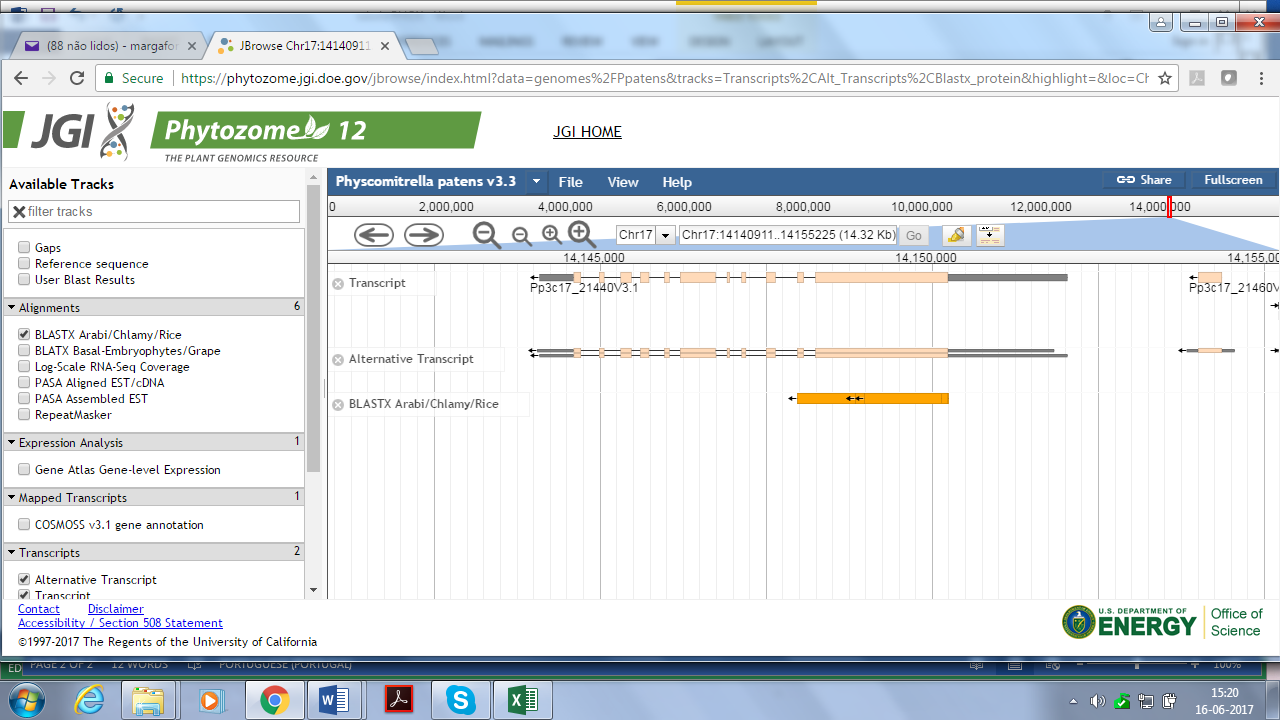

Pp3c2_2780

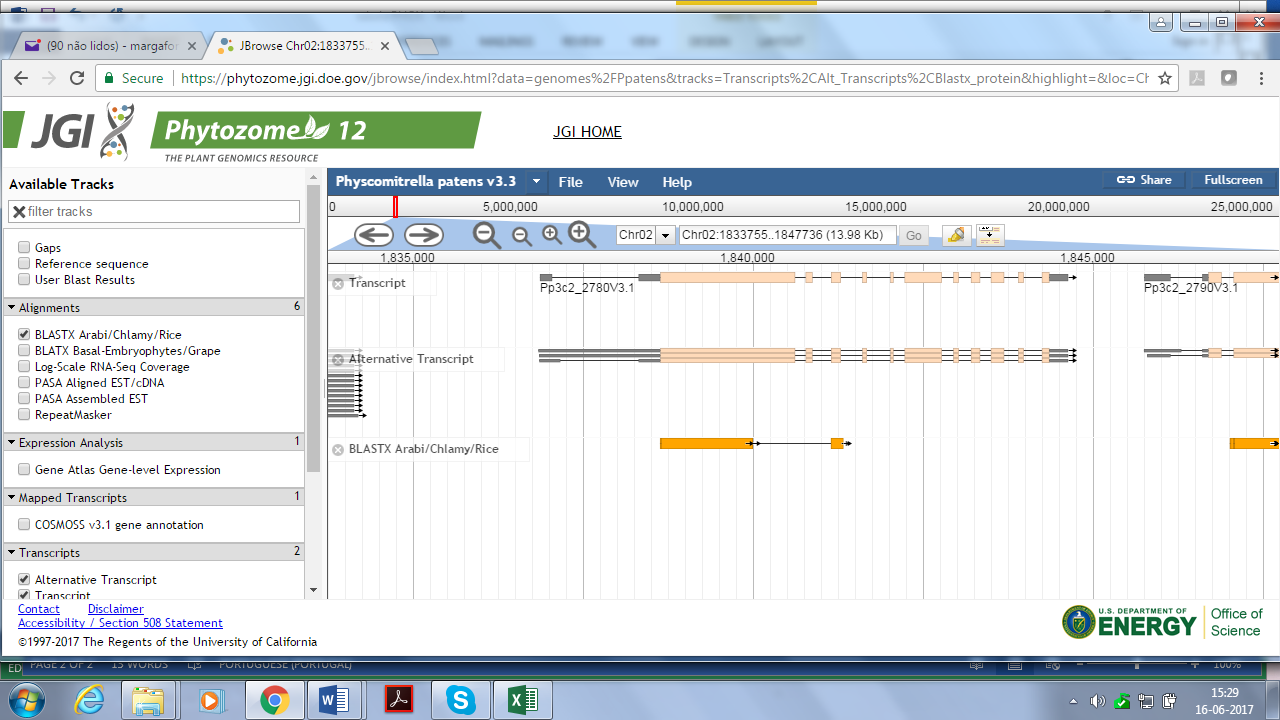

Pp3c14_16910

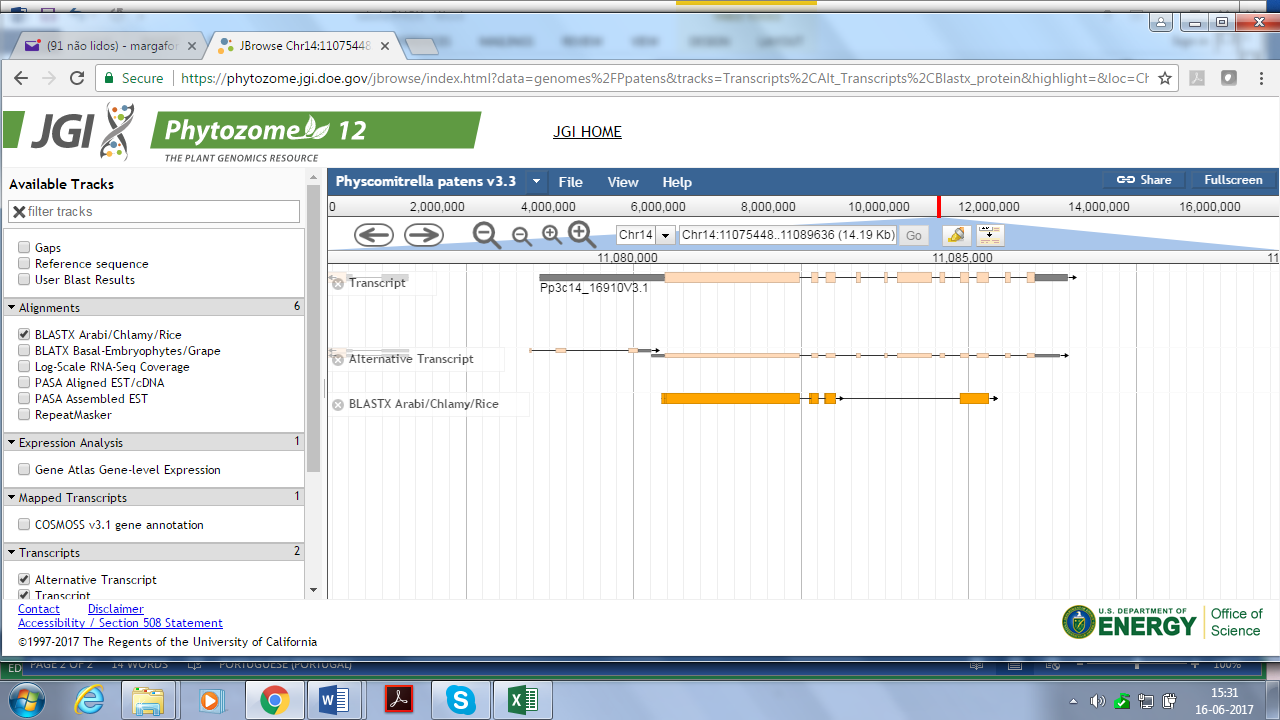

Pp3c10_18970

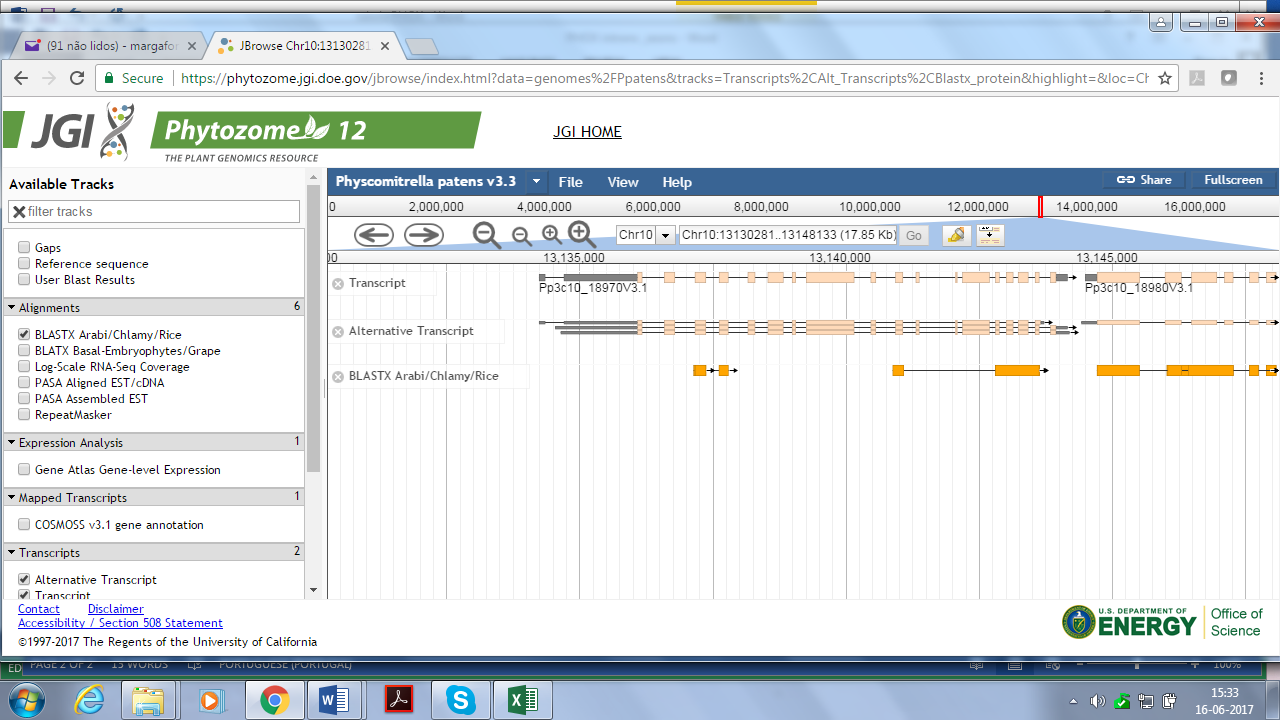

**CLASS IV**

**Pp3c12_3620**

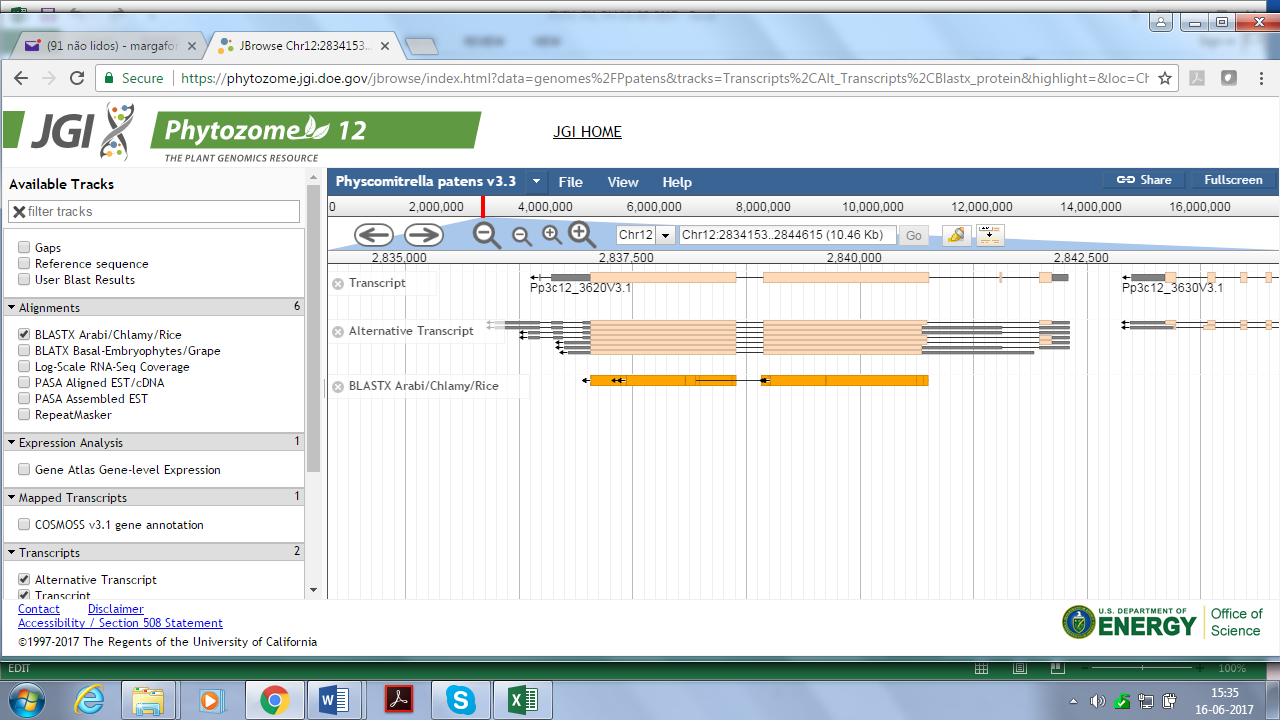

Pp3c13_3440

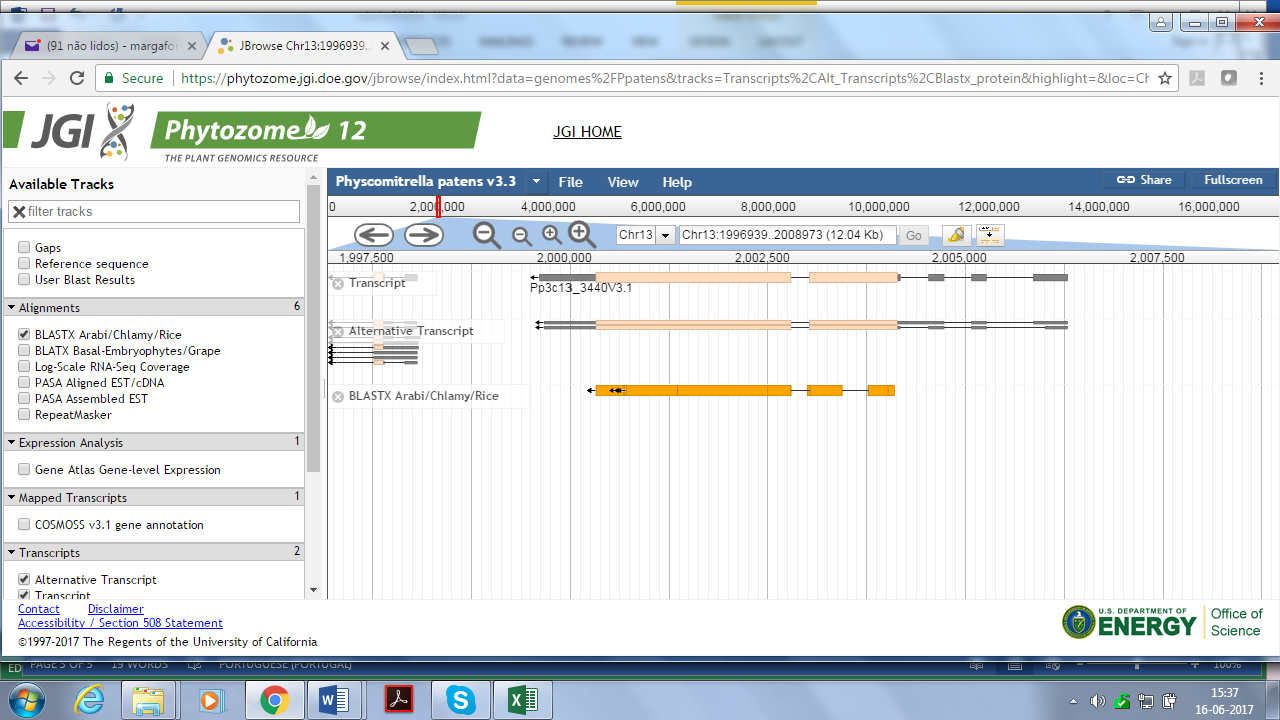
